# Epithelial organ shape is generated by patterned actomyosin contractility and maintained by the extracellular matrix

**DOI:** 10.1101/2020.01.22.915272

**Authors:** Ali Nematbakhsh, Megan Levis, Nilay Kumar, Weitao Chen, Jeremiah Zartman, Mark Alber

**Affiliations:** Department of Mathematics, University of California, Riverside, CA; Interdisciplinary Center for Quantitative Modeling in Biology, University of California, Riverside, CA; Department of Chemical and Biomolecular Engineering, University of Notre Dame, Notre Dame, IN; Bioengineering Graduate Program, University of Notre Dame, Notre Dame, IN; School of Medicine, University of California, Riverside, CA; Department of Bioengineering, University of California, Riverside, CA

**Keywords:** morphogenesis, cell mechanics, multiscale, computational model, *Drosophila* wing disc

## Abstract

Epithelial sheets play important roles in defining organ architecture during development. Here, we employed an iterative experimental and multi-scale computational modeling approach to decouple direct and indirect effects of actomyosin-generated forces, nuclear positioning, extracellular matrix (ECM), and cell-cell adhesion in shaping *Drosophila* wing imaginal discs, a powerful system for elucidating general principles of epithelial morphogenesis. Basally generated actomyosin forces are found to regulate apically biased nuclear positioning and are required for generating epithelial bending and cell elongation of the wing disc pouch. Surprisingly, however, short-term pharmacological inhibition of ROCK-driven actomyosin contractility does not impact the maintenance of tissue height or curved shape. In comparison, the relative tautness of the extracellular basement membrane is also patterned between regions of the wing disc. However, computational simulations show that patterning of ECM tautness provides only a minor contribution to modulating tissue shape. Instead, the buildup of a passive ECM pre-strain serves a principle role in shape maintenance. Surprisingly, this is independent from the maintenance of actomyosin contractility. Furthermore, localized apical adhesion between the two cell layers within the wing disc requires ROCK-driven actomyosin activity in the absence of the basal extracellular matrix. This apical adhesion between the two cell layers provides additional mechanical support to help maintain tissue integrity. The combined experimental and computational approach provides general insight into how the subcellular forces are generated and maintained within individual cells to induce tissue curvature and suggests an important design principle of epithelial organogenesis whereby forces generated by actomyosin followed by maintenance as pre-strain within the ECM are interconnected, but functionally separable.

**Significance statement:** A major outstanding question in developmental biology is the elucidation of general principles of organ shape formation and maintenance. Here, an iterative experimental and multi-scale computational modeling approach reveals that actomyosin contractility generates the bent profile along the anterior-posterior axis while tension within the ECM is sufficient and necessary for preserving the bent shape even in the absence of continued actomyosin contractility once the shape is generated. The mechanisms tested in this study define the necessary factors for establishing the shape of the wing disc, which later everts to form the adult wing during pupal development. The method can be extended to test novel mechanisms of other epithelial systems that consist of several cellular and ECM layers.

## Main Text

### Introduction

Epithelial tissues are critical drivers of morphogenesis^1–3^. Functionally, they serve as barriers between the environment and internal structures of organs. Bending and folding are common features of many epithelial tissues^4^. However, a predictive understanding of how organs regulate their shape at a given stage of the development remains elusive. This is partially because the roles of mechanical properties of components of cells and tissues during organ development are hard to quantify experimentally. Further, the interactions between subcellular components that define tissue level-properties are non-linear, non-intuitive and time-varying. Elucidating general design principles that can explain the overall mechanisms governing epithelial morphogenesis remains a key goal for characterizing multicellular systems^5–7^. Consequently, computational modeling approaches coupled to experimental studies are becoming powerful new tools to infer and test the basic design principles of epithelial morphogenesis.

The *Drosophila* (fruit fly) wing imaginal disc serves as a paradigm system to study epithelial morphogenesis (Fig. 1)^8–10^. A genetic and biophysical toolkit that includes recent advances in organ culture and live-imaging techniques has been developed to investigate mechanisms underlying the shape formation of a wing disc^6, 7^. During larval stages (1^st^, 2^nd^, and 3^rd^ instar), the wing disc undergoes a period of rapid growth with significant shape changes from a round epithelial vesicle consisting of a single epithelial monolayer^10, 11^. At early stages of development, the wing disc, consisting of cuboidal cells, develops into a stereotypically folded tissue with multiple classes of epithelial cells, including squamous, cuboidal and pseudostratified columnar cells^12^. In mid-to late larval stages, the wing pouch forms multiple folds along the dorsal-ventral axis while a characteristic bent “dome” shape in the cross-sectional profile along the anterior-posterior axis is stably maintained (Fig. 1C-E, SI Fig.1)^13–15^. The stereotypical shape of the wing disc plays an important role as the initial geometric condition leading into pupal morphogenesis. During pupal morphogenesis, the wing disc undergoes a series of morphogenetic steps to form the adult fly wing^10, 16, 17^. A failure to achieve and maintain this stereotypical folded and curved shape at the end of larval development distorts subsequent stages of wing disc morphogenesis, resulting in misshapen wings. In turn, the final wing shape is critical for ensuring adequate flight performance^18^.

**Figure 1.**
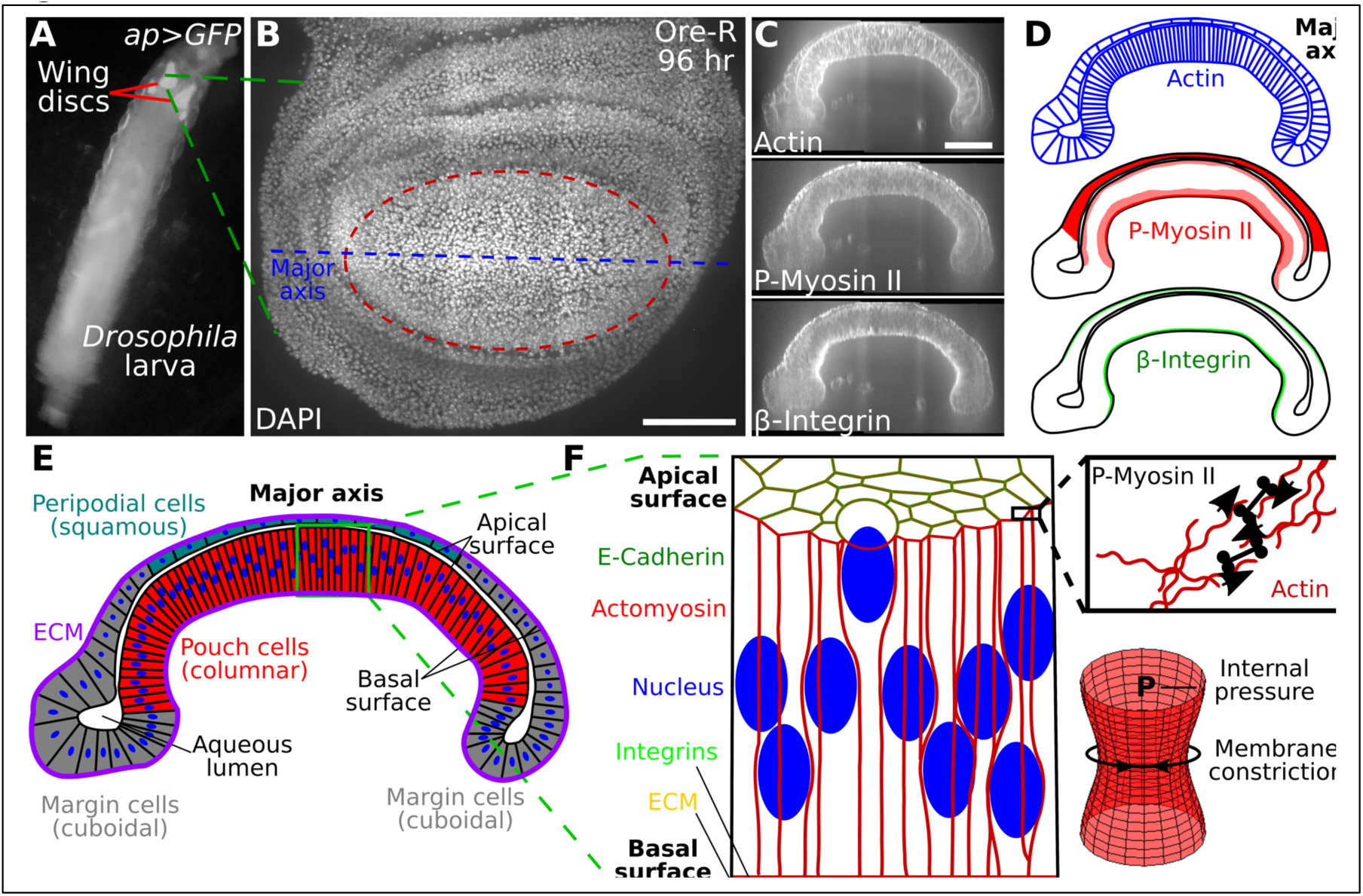
A) Imaginal wings discs in 3^rd^ instar *ap>GFP* larvae. B) Top view of a wing disc at 96 hours after wing disc in B stained for actin (phalloidin), P-Myosin II, and β-Integrin. C, D) Actin, P-Myosin II and β-Integrin patterns in the major axis cross-section. E) Multiple cell types within the major axis cross-section. The wing disc is composed of squamous peripodial cells (top, blue), columnar pouch cells (bottom center, red), and cuboidal marginal cells (sides, grey). An aqueous lumen is enclosed by the apical surface of the two cell layers. The basal surface is constrained by the extracellular matrix (ECM). F) Structural components of the *Drosophila* wing disc columnar cells (left). Schematic showing for actomyosin contractility (right, top). An actin mesh provides structural support to the cell membrane. Actin filaments are pulled together by phosphorylated non-muscle Myosin II (P-Myosin II) resulting in increased membrane tension. Actomyosin driven tension opposes the internal pressure in cells and acts to constrict the membrane (right, bottom). Scale bars are 50 μ*m*.

Key structural proteins including actin, phosphorylated non-muscle Myosin II (P-Myosin II), and β-integrin are patterned within the larval wing disc (Fig. 1C, D). The cross-section of a 96 h after egg laying (AEL) wing disc consists of squamous (peripodial) and columnar (pouch) cell layers that adhere to each other along their apical surfaces. Connected together, these cells enclose an aqueous lumen^19^. A thin extracellular matrix (ECM) forms a basement membrane that wraps around the cells along their basal surface (Fig. 1E, F). Integrins adhere the ECM to the basal surface of cells while the integrity of the apical surface is maintained by intercellular adhesion junctions (Fig. 1F). An actin mesh containing P-Myosin II (actomyosin) provides structural integrity to individual cell membranes (Fig. 1F). Actomyosin promotes membrane constriction and thus opposes internal cellular pressure.

Folding mechanisms have been widely explored for multiple tissues, including the developing wing disc^20–31^. However, how subcellular processes contribute to the global shape of the tissue is a fundamental, unanswered question. In this study, we investigated the contributions of multiple cellular processes to the regulation of the overall stereotypical dome shape along the major anterior-posterior (AP) axis of the wing disc. To do so, we combined quantitative experimental analysis with a newly developed and biologically calibrated, subcellular element-based computational model. In the computational model, we included cell membrane, nuclear shape and position, and homotypic and heterotypic cell-cell and cell-ECM interactions. In the experimental analysis, we measured multiple features of wing discs at 96 h AEL, including nuclear positioning, cell height and tissue shape in the wild type condition. We also imaged the spatial distribution of actomyosin, collagen and integrins. These experimental data provided important inputs to calibrate the computational model.

Results from computational model simulations and quantitative analysis of genetic and pharmacological perturbations of the wing disc were integrated to explain how subcellular dynamics impact the formation of the tissue level features of the wing. We found that patterned actomyosin contractility explains the apically biased nuclear positioning and the global bending shape generation along the AP axis observed in experiments. Surprisingly, actomyosin contractility is not required for maintaining the bending shape. While patterned pre-straining of the underlying ECM only plays a minor role in generating the curved tissue shape along the AP axis of the organ, it is critical for maintaining that bent shape once generated. Lastly, perturbation studies demonstrated that maintaining regional apical adhesion between the two cell layers within the wing disc requires ROCK-driven actomyosin activity in the absence of the basal extracellular matrix.

## Results

### The distribution of actin, phosphorylated myosin II and β-integrin is patterned along the wing disc cross-section

Quantitative spatiotemporal maps of subcellular components are required to develop integrative models of organogenesis. We first quantitatively mapped the primary structural components involved in regulating mechanical properties of cells in 3^rd^ instar wing discs. Discs were obtained at multiple stages and stained for key architectural components of cells (Fig. 1, 2 and SI Fig.1). To our knowledge, the biophysical mechanisms determining the overall cross-sectional profile along the AP axis have not been quantitatively investigated. Visualization of fluorescent intensities along the major axis in this cross-section showed that filamentous actin (F-actin) was consistently localized to cell membranes (Fig. 1C, D). Phosphorylated non-muscle Myosin II (P-Myosin II) was seen in squamous cells and at higher intensity levels in columnar pouch cells within a narrow strip along the apical surface and a wider strip between the basal surface and nuclei (SI Fig.1, 5). β-integrin was concentrated basally in columnar cells and to a lesser degree in squamous cells (Fig. 1C, D and SI Fig.1, 5). These protein distributions suggest actomyosin-driven activity is significant in squamous cells and the apical surface and basal compartment beneath cell nuclei in columnar cells. Observed cell shapes suggest that this contractility occurs laterally in squamous cells resulting in flatter cells and apicobasally in columnar pouch cells, such that pouch cells become narrower and more elongated. The differences in β-integrin intensity associated with squamous and columnar cells suggest differential relative integrin-ECM adhesion between the two layers and possible heterogeneity in passive tension levels in the underlying extracellular matrix, which requires additional analysis.

**Figure 2.**
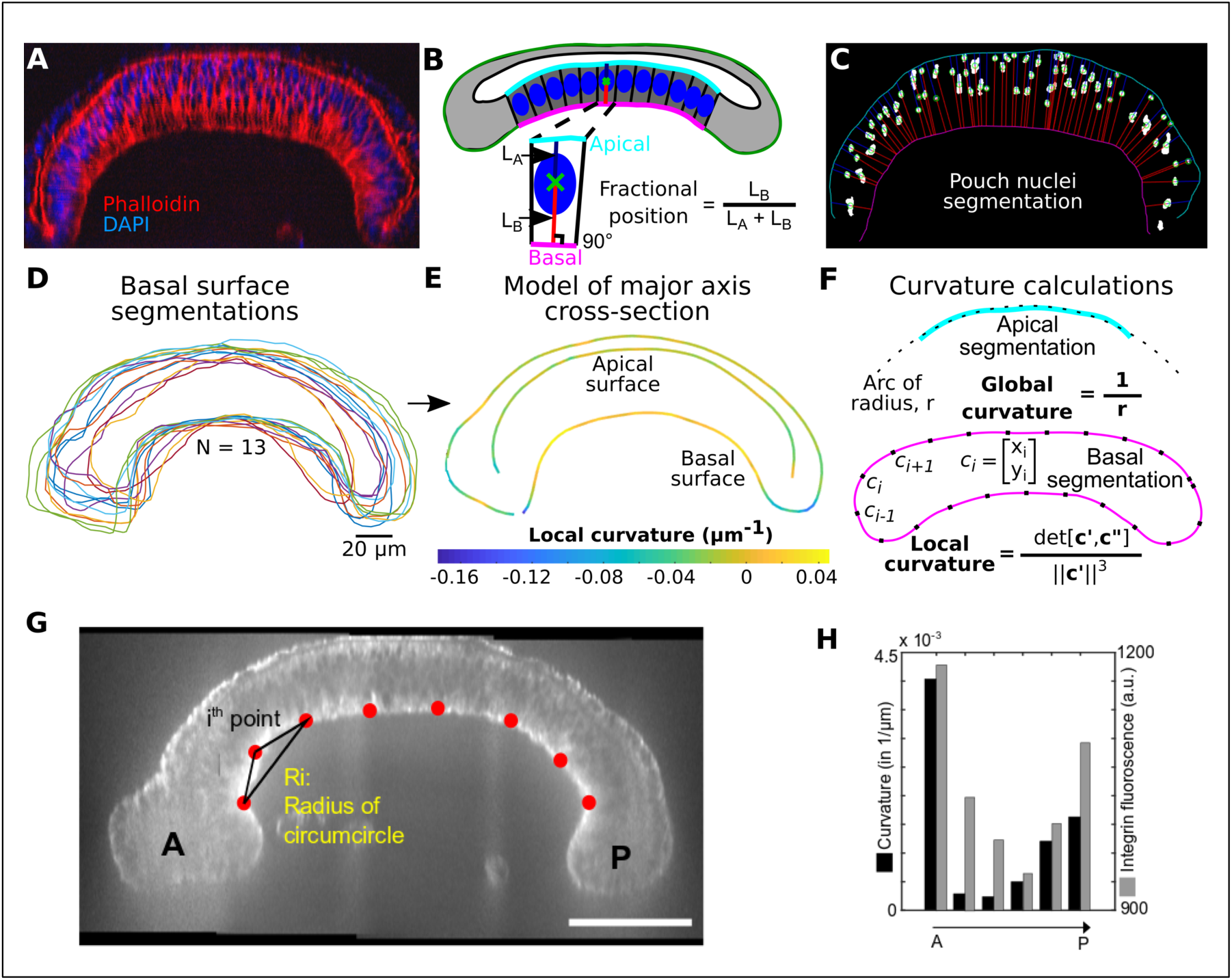
Quantitative analysis of wing disc shapes and nuclear positions along the major axis of imaginal wing discs at 96 hours AEL. A) Representative pre-processed image of a wing disc cross-section stained with DAPI (nuclei) and phalloidin (staining F-actin). These images were used for nuclei and surface segmentation. B) The fractional position between the apical and basal surface serves as a metric for quantifying relative nuclear positions. Note: heights of columnar pouch cells were calculated as the sum of LA and LB. C) Representative segmentation of nuclei. Blue and red lines correspond to LA and LB in B, respectively. D) Basal surface segmentation for multiple samples in different colors (N=13). E) Composite representation of wing disc morphology generated from samples in D. F) Quantification of global and local curvatures. Supplemental SI Fig. 2 and SI Fig. 3 provide additional details on the image processing pipeline. G) Representative image highlighting average curvature quantification used for H, the curvature was found using a circumcircle approach. (Scale bar is 50 μ*m*). H) Curvature quantification of regions of interest along the pouch as well as Intensity quantification of integrin along the basal side of the pouch.

Additionally, fluorescently stained nuclei exhibit apically-biased positions in columnar pouch cells (Fig. 2A). We segmented nuclei and the apical and basal surfaces of the pouch region to quantify the fractional apicobasal nuclear position (Fig. 2B, C). This revealed that nuclear positions in columnar cells are biased toward the apical surface (SI Fig. 2 and SI Fig. 3). Moreover, this nuclear migration in wing discs depends on actomyosin contractility^32^. Since the actomyosin is present at the basal side of the columnar cells, we hypothesized that actomyosin may also contribute to the tissue-level dome, or bent, shape of wing disc along AP axis.

### The curvature of the basal surface correlates with local β-integrin concentration levels

Images of wing discs stained for actin and nuclei were processed to extract apical and basal surface segmentations (Fig. 2D). The extracted profile measured by the segmentation framework (SI Fig. 2-4) provides generalized shape information of the wing disc cross-section, including tissue thickness and local curvature measured for the composite pattern of the basal surface for experimentally captured tissues (Fig. 2E). A comparison of this local curvature profile to the distribution of β-integrin (Fig. 2G, 2H) reveals a correlation between basal curvature and β-integrin intensity. Marginal cells at the lateral edge of the tissue (here termed boundary cells) had a reduced signal of β-integrin accumulation and exhibited a strongly negative basal surface curvature. The basal surface associated with squamous cells had a higher amount of β-integrin localization than boundary cells and slightly negative basal curvature. In the pouch region, β-integrin density was higher than both boundary cells and squamous cells, and the corresponding basal curvature was positive.

These observations suggest that β-integrin-associated adhesion results from an accumulation of passive ECM tension. Imaging of wing discs with fluorescently labeled Collagen IV (Viking::GFP) revealed that the ECM was relatively loose around the peripodium. In contrast, it is taut along the basal side of the pouch (Fig. 3A), further supporting a possible role for β-integrin-mediated regulation of ECM tension.

**Figure 3.**
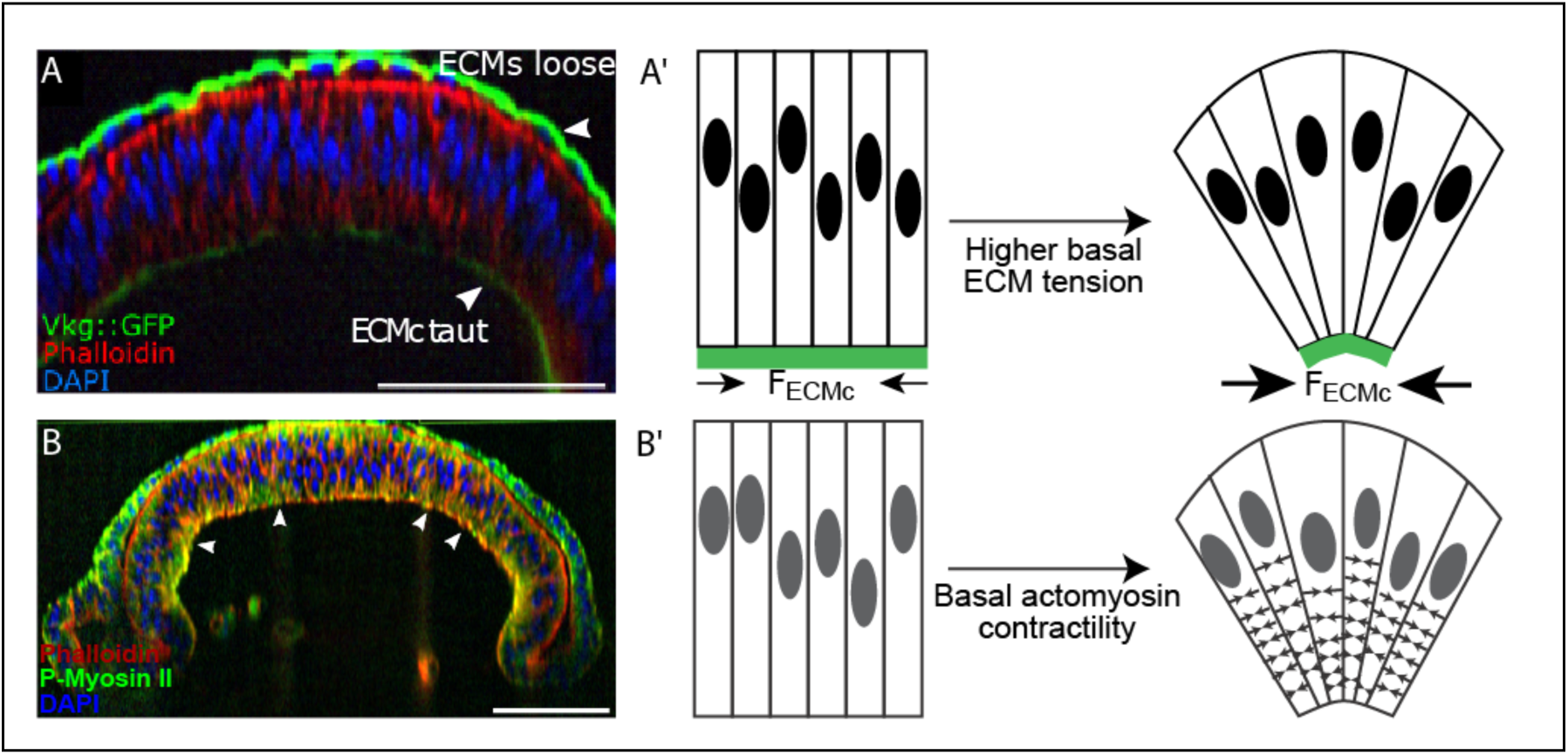
Potential mechanisms of wing disc bending: A) The patterned ECM tension hypothesis. In this hypothesis, the level of passive tension of the ECM is higher next to the columnar cells compared with ECM next to squamous cells. Scale bar is 50 μ*m*. A’) Graphical illustration of potential hypothesis explaining how high ECM tension at the basal side of the columnar cells compresses the tissue and contributes to curvature profile of wing disc. B) The patterned actomyosin contractility hypothesis. The columnar cells were stained for actin by Phalloidin and an antibody to P-Myosin II (Scale bar is 50 μ*m*). B’) Schematic of hypothesized mechanism for generating epithelial bending and asymmetrical nuclear positioning in the wing imaginal disc. Actomyosin contractility beneath the nucleus of columnar cells drives bending of the wing disc.

### Experimental observations require a multiscale mechanical model to determine mechanisms of relative contributions of subcellular components to overall tissue shape

The apically biased nuclear positioning in the pouch region suggests a possible mechanism for bending the wing disc along the anterior-posterior axis (SI Fig. 3). Consistent with nuclei being an order of magnitude stiffer than cytoplasm^33^, columnar cells are wider near the apical surface. This would, in turn, tend to result in stretching and bending forces on the squamous cells that consist define the top layer of the wing disc. Further, the apical asymmetry in the nuclear position possibly facilitates bending of the tissue (Fig. 2). Thus, we hypothesized that actomyosin-driven forces might play a dominant role in maintaining all nuclei near the apical side in columnar cells, also resulting in the bending shape of the epithelial monolayer (Fig. 3B-B’). On the other hand, experiments also suggest that the ECM provides lateral compression to the wing disc (Fig. 3A-A’) as the columnar pouch cells are flatten out and become more cuboidal upon chemical digestion of the ECM with collagenase^34, 35^. However, whether passive ECM tensile forces or active actomyosin contractility can individually account for either generating or maintaining the bent shape along the anterior-posterior axis remains unclear^15^.

Disentangling the interplay between subcellular architectural components including actomyosin, integrin, and extracellular matrix and their associated mechanical forces is experimentally challenging. Thus, we created a combined computational model and experimental approach described in the Method section to test proposed mechanisms individually. We formulated a subcellular element model that enables us to simulate the subcellular components of cell architecture in details (Methods section and Fig. 10). Application of a detailed multiscale model in two dimensions reasonably describes the essential geometric features of wing disc pouch. In particular, the curved shape along the major pouch axis (AP Axis, Fig. 1B, SI Fig. 1) is representative of the characteristic “dome” shape throughout the pouch region.

**Figure 10.**
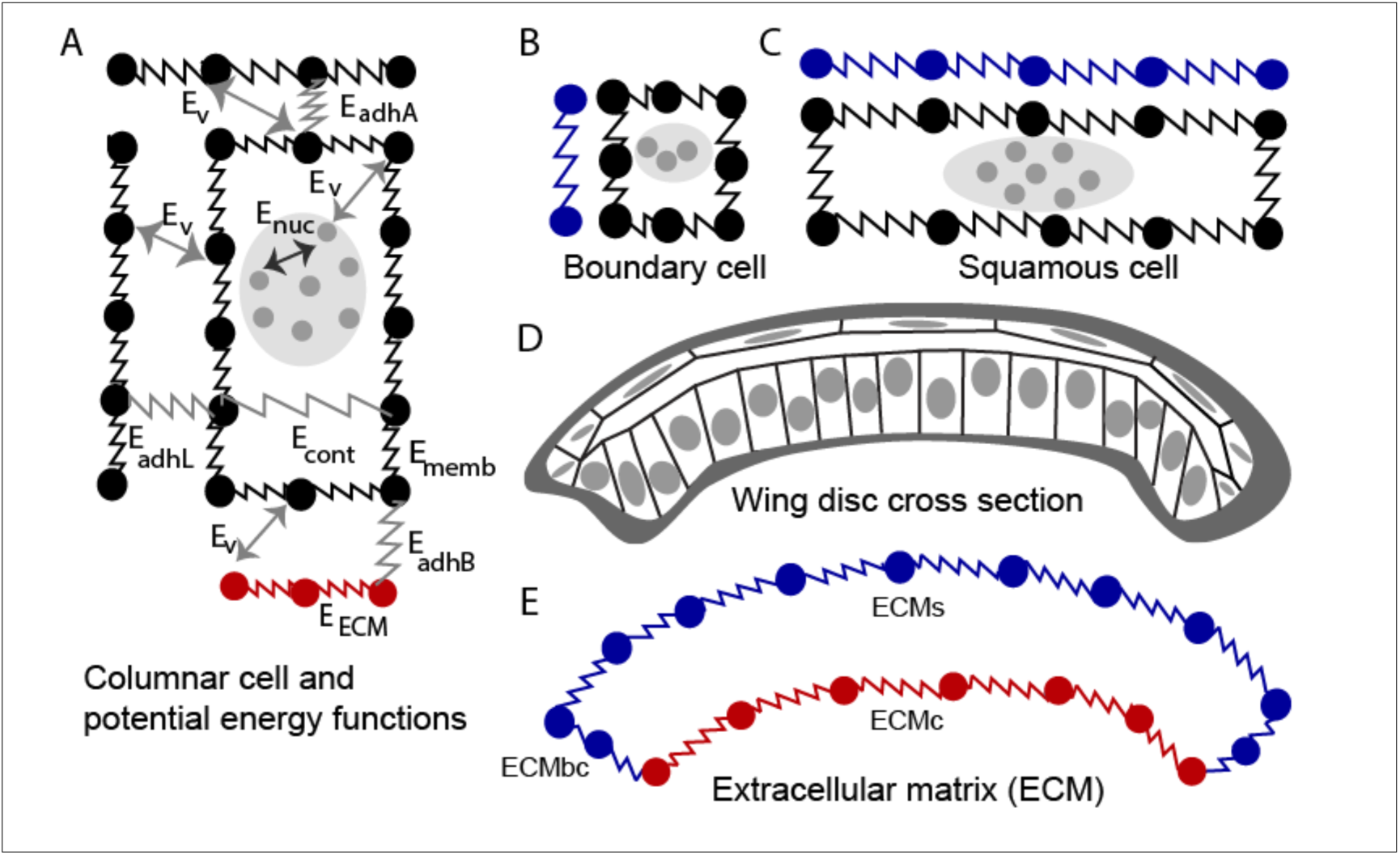
Diagram of the subcellular element (SCE) model of the cross-sectional profile of the wing along the anterior-posterior axis. A) Columnar cell submodel with its potential energy functions describing interactions between adjacent cells, intracellular interactions and cell-ESM interactions. B) Marginal boundary cell submodel. C) Squamous cell submodel. D) Diagram of the cross-sectional profile of the wing along the anterior-posterior axis which includes columnar cells, boundary cells, squamous cells and ECM. E) Diagram of the submodel of the ECM divided into separate sections: ECMc, ECM associated with columnar cells in the wing pouch; ECMbc, ECM associated with the marginal boundary cells at the lateral region of the disc, and ECMs, ECM associated with squamous cells.

We incorporated into the computational model mechanical spring-like forces generated by actomyosin at the subcellular level, passive tension within the ECM, and adhesion between two cell layers consistent with previous observations^19, 36, 37^ to investigate their individual functional roles. Composite experimental profiles used throughout this study provided both qualitative and quantitative comparisons with predictions from computational simulations.

### Passive tension within the ECM is not sufficient to bend the tissue

Recent work has demonstrated that the ECM actively contributed to the morphology of epithelial cells in the pouch^34, 38^. In particular, removal of the ECM reduced lateral compressive forces on the columnar cells in the *Drosophila* wing disc and leads to shorter and fatter cells (SI Fig. 4). Therefore, these previous results suggest that the ECM is under tensile stress and it in turn compresses the cells. Consequently, we investigated whether the passive tensile stresses within the ECM was necessary and sufficient to generate epithelial bending of the wing disc.

As previously mentioned, the imaginal wing disc is enclosed by the ECM, which connects to the basal side of individual cells through integrin-mediated adhesion^38^. ECM remodeling follows the growth and division of epithelial cells in the wing disc^12^. A high growth rate of epithelial cells in comparison with ECM remodeling, can lead to accumulation of tensile stress in the ECM^39^. In other contexts, cell stresses are converted into pre-strains within the ECM^40^. Cell division happens more frequently in the columnar pouch cells than in the squamous peripodial cells^12^. This led us to assume that the tensile stress is higher in the basement membrane associated with the columnar cells. Experimental images confirmed that assumption as the ECM is relatively loose around the peripodium whereas it appeared taut along the pouch (Fig. 3A, B).

Additionally, there is significantly higher observed integrin intensity connecting the basal side of cells to ECMc (Fig. 1C). We first tested the hypothesis that higher tensile stress in ECMc in comparison with ECMs can lead to the overall bending shape in *Drosophila* wing disc (Fig. 3A’). To test this, we performed computational model simulations without accumulated tension in the ECM (Fig. 4B-D, Baseline) as a baseline reference case to determine the shape of the tissue without a mechanism of active force generation. This baseline case was then compared to simulations with increased levels of tensile stress accumulated in the ECMc. Then we tested a model scenario with uniform tensile stress in both the ECMc and ECMs. Finally, we tested cases with differential stresses between each side of the tissue. As absolute levels were unknown, we varied the ECMc tensile stress from 4 to 7-fold higher than the tensile stress of the ECMs. All simulations start with a flat tissue and asymmetric nuclear distribution (Fig. 4A) as the initial condition of the tissue morphology. Each simulation was run until a constant steady state configuration of tissue was reached (Fig. 4A’).

**Figure 4.**
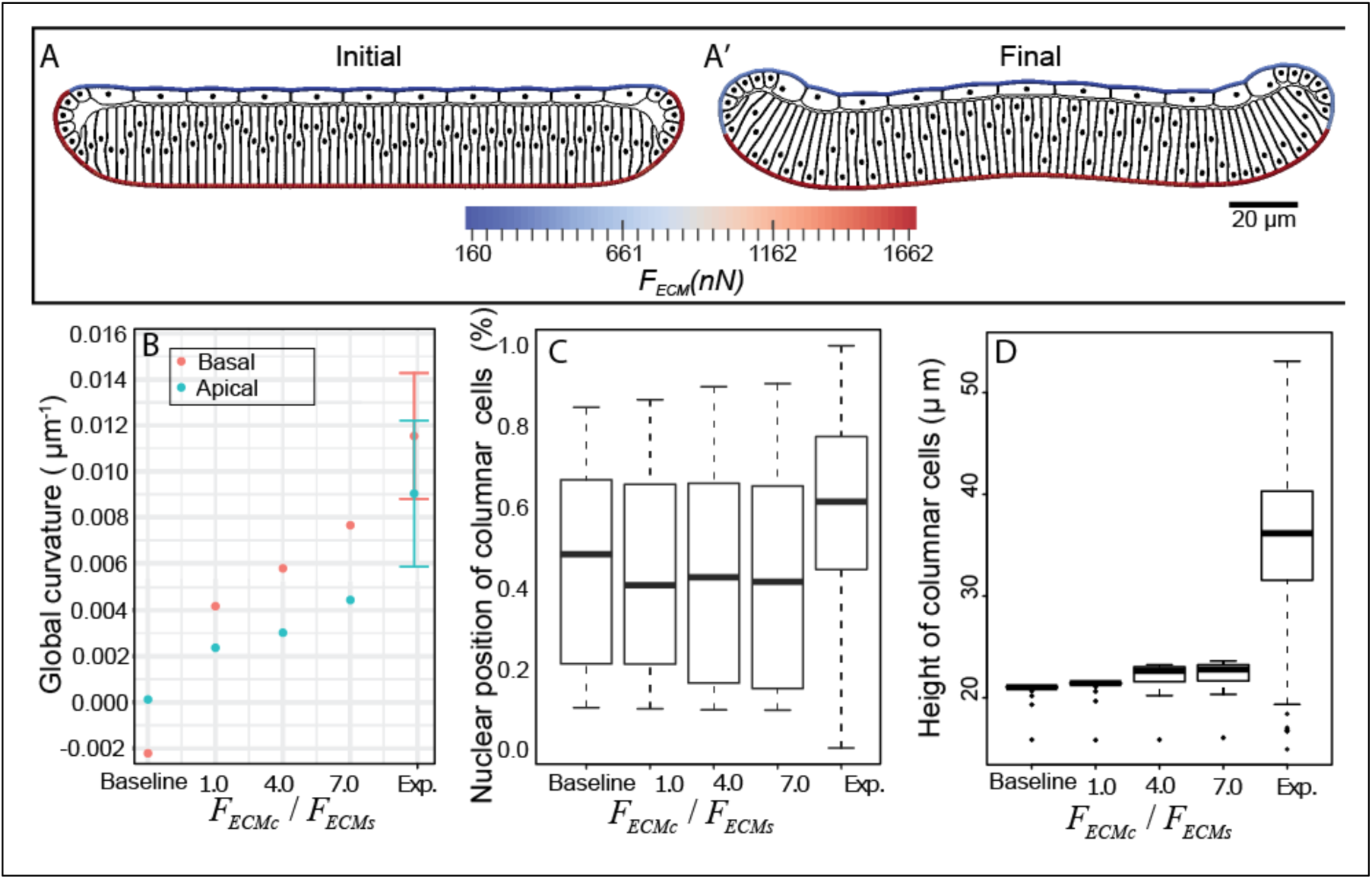
Computationally testing the hypothesis of patterned ECM tension reveals that direct ECM tension cannot generate epithelial bending. In these simulations, the difference between the passive tensile stress of the ECM is associated with squamous cells (F_ECMS_) is lower than the passive tensile stress of the ECM associated with columnar cells (F_ECMc_ ). Such a difference is insufficient to explain the bending of the wing disc pouch, but it contributes to the overall architecture of the wing disc. A) Initial and final frames of a representative computational simulation showing how higher tension in ECM associated with columnar cells in comparison with the tension in the ECM associated with squamous cells leads to curved shape profile of the wing disc. Comparison of in-silico prediction of impacts of ECM level of tension on B) curvature profile of experimental (n=16) and simulated (n=1) wing discs. Experimental data are mean ± standard deviation. Simulation result of final global curvature of the tissue is unique for every set of input parameters, so that one simulation is sufficient. C) relative position of nuclei, D) height of columnar cells. In C and D, boxplots show minimum, first quartile, mean, third quartile, maximum and outlier of simulated (n=65) and experimental (n=1064) columnar cells. “Baseline,” here, corresponds to the condition where the whole ECM initiated without any tension.

A comparison of simulated and experimental results in Fig. 4B demonstrates that curvature profiles of the top and bottom surfaces of the simulated wing disc in all the cases are significantly less than the curvature profiles obtained in corresponding experiments. Moreover, nuclear positions equilibrated toward a symmetrical distribution along the apical-basal axis of columnar cells for all sets of simulations. Moreover, the final average height of columnar cells shown in Fig. 4D is considerably less than 25 μm, which is inconsistent with the experimental observations. The overall results indicate that the tensile stress within ECM contributes little to the overall bending shape of the tissue and cannot explain the asymmetric nuclear distribution within the tissue.

### Basal actomyosin contractility is sufficient to induce tissue bending

Actomyosin plays an important role in shape formation during cell growth and tissue development^15, 16, 32^. For example, actomyosin contractility drives the nuclear motion in the columnar cells of *Drosophila* wing discs to enlarge the apical side during mitosis^32^. We hypothesized that actomyosin contraction beneath the nuclei of columnar cells not only maintains the asymmetrical distribution of nuclei along the apical-basal axis, but also bends the wing disc tissue (Fig. 3B-B’). Therefore, we performed a set of simulations with different levels of basal actomyosin contractility to test the global effect of basal actomyosin contractility beneath nuclei in columnar cells on tissue shape. To ensure we are only observing the effect of basal actomyosin contractility, we did not include any ECM tensile force at the initial time point.

These simulations started with a flat tissue and asymmetric nuclear distribution, while the basal sides of pouch cells contracted as shown in Fig. 5A. The simulations led to nuclear motion, changes in cell size and shapes, and tissue bending until the simulation reaches a final steady state. Three sets of simulations were performed with low (3 *nN*/μ*m*), medium (6 *nN*/μ*m*) and high (9 *nN*/μ*m*) levels of actomyosin contractility. Each condition was then compared with both experimental data and the baseline simulations where there is no basal actomyosin contractility.

**Figure 5.**
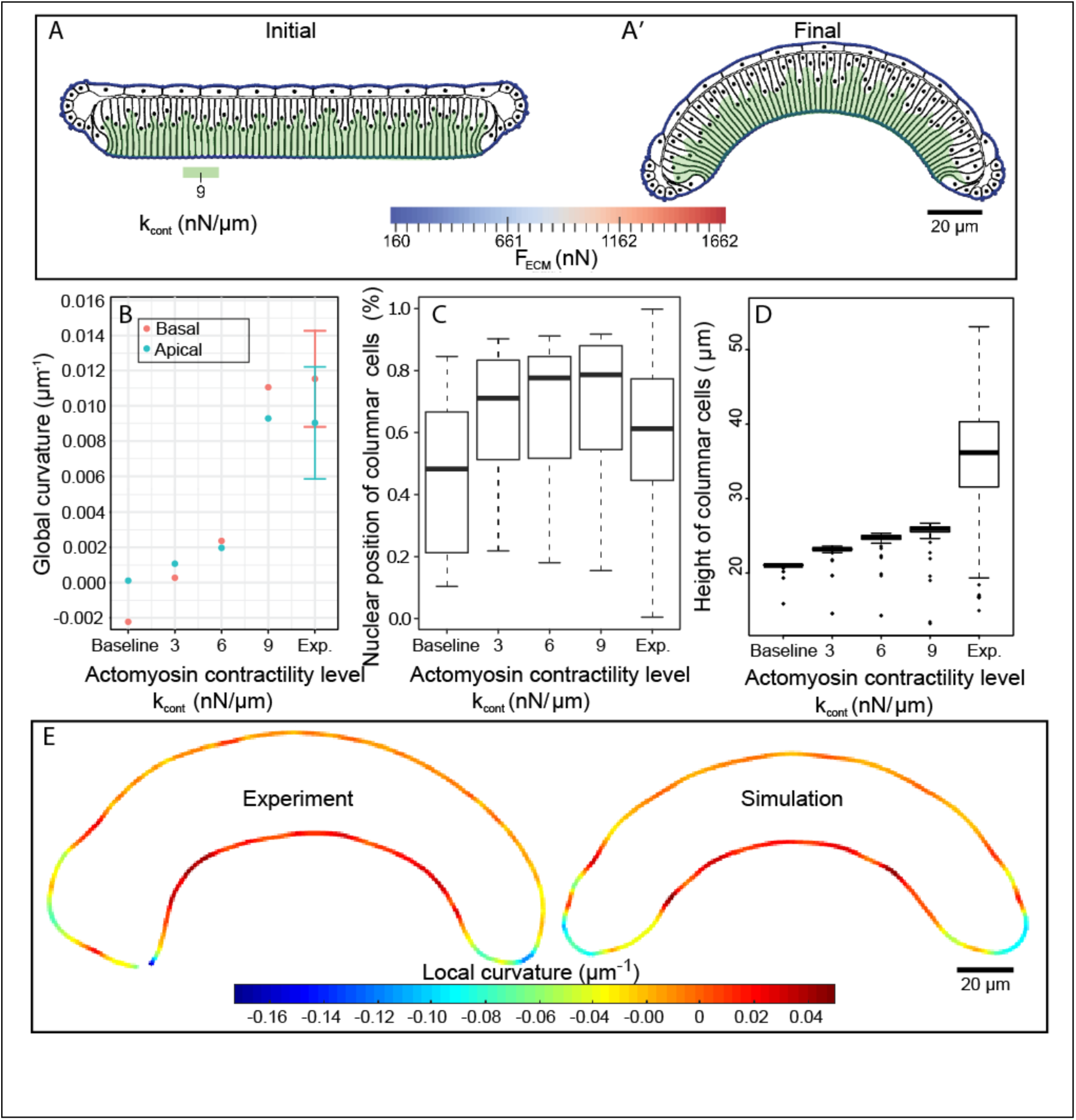
Quantitative and qualitative comparisons of experimental data with simulation results for different levels of simulated basal actomyosin contractility (k_contract_). A) Initial and final frames of simulation result where actomyosin contractility leads to curved profile shape. B) Global curvature of apical pouch surface and basal peripodial surface for experimental (n=16) and simulated (n=1) wing discs. Experimental data are mean ± standard deviation. C) Relative nuclei positions of pouch cells. Note, the relative nuclei position has a value of zero at the basal surface and one at the apical surface. D) Height of pouch cells. In C and D, boxplots show minimum, first quantile, mean, third quartile, maximum and outlier of simulated (n=65) and experimental (n=1064) columnar cells. Measurements for columnar cells height, experimentally and computationally were taken as the summation of the length of straight lines from apical to centers of segmented nuclei centers and from segmented nuclei centers to the basal side of the cells. E) Comparison of local curvature profiles and shapes obtained from experimental data and computational results.

Simulations with different levels of basal actomyosin contractility demonstrated a reduction in the width beneath the nuclei of the columnar cells, resulting in the overall wing disc bending in the same direction as observed in experiments (Fig. 5A’). The tissue curvature increased as a function of the level of actomyosin contractility (Fig. 5B). Moreover, simulations demonstrated that the apically-biased nuclear distribution was maintained in columnar cells, even with minimal levels of actomyosin contractility (Fig. 5C). Heights of columnar cells also increased with increased contractility levels (Fig. 5D), although it is still less than experimental data. A plausible reason for a reduced height of cells in simulations compared with experimental data is due to initial height of columnar cells in the simulation where lower range of experimental data are chosen for computational cost efficiency. The local curvature profile along obtained in model simulations with high contractility is in a very good agreement with the profile observed in experiments (Fig. 5E).

Application of the sensitivity analysis method based on Latin Hypercube sampling and partial correlation coefficient method ^41^ (Section SI.5) confirmed that the overall shape (curvature) is primarily generated by patterned actomyosin contractility, with higher contractility below nuclei. This computational result confirms our hypothesis that the basal actomyosin contractility plays a significant role in bending the tissue along the AP axis. Model simulations predict that the actomyosin generates 9 *nN*/μ*m* of force to produce the experimentally observed bent shape. Therefore, actomyosin contractility beneath nuclei induces the asymmetric spatial distribution of nuclei in columnar cells and curved shape for the entire tissue that are observed experimentally. This suggests an effective mechanism for generating the curved shape of the *Drosophila* wing along the anterior-posterior axis.

To test the main prediction that active actomyosin contractility is essential for generating the curved profile of the wing disc along the anterior-posterior axis with cell height and nuclear position controlled, we investigated the impact of knocking down Rho1, a key egulator of actomyosin contractility (Fig. 6). We used the GAL4/UAS binary expression system which allows for tissue-specific expression of RNAi constructs. The MS1096-Gal4 driver has higher expression in the dorsal compartment and limited expression in the ventral compartment. Based on previous research, we used an RNAi line that does not lead to qualitative phenotypic changes as a control (Ryr^RNAi^ expression by the same Gal4 driver). Columnar cell height was quantified by calculating average cell heights in the anterior and posterior side of the wing disc as shown in B”. Inhibition of Rho1 in the wing imaginal discs lead to a significant increase in columnar cell height (Fig. 6E), in agreement with previous findings for inhibiting Rho1^13^. Further, bending was quantified by calculating the local (Manger) curvature of the basal surface (Fig. 6F). In agreement with computational model prediction, the inhibition of Rho1 in the wing imaginal discs resulted in a significant reduction in tissue curvature (p-value from t-test ∼10^-5^).

**Figure 6:**
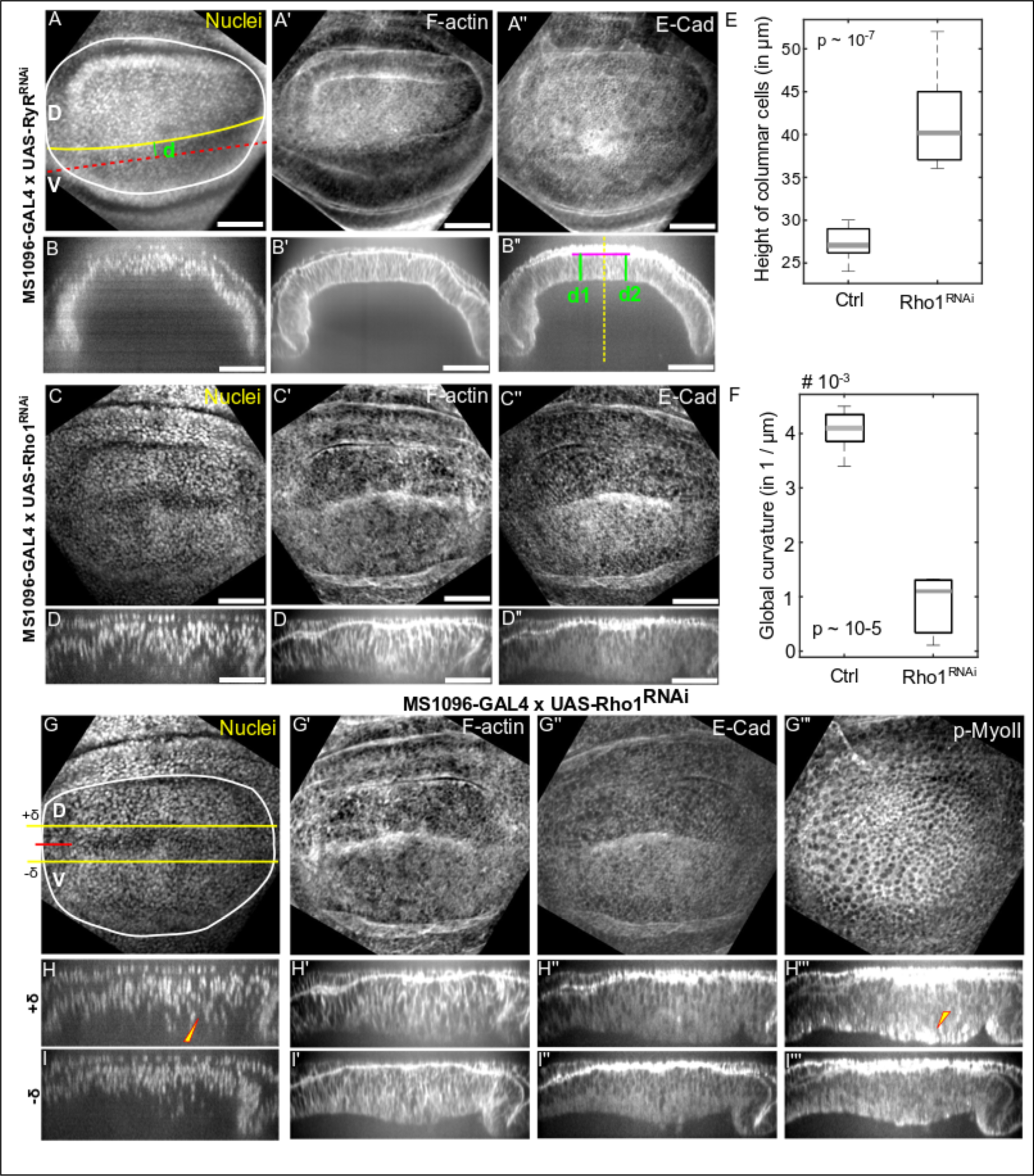
Rho1 promotes tissue bending and regulation of cell height. (A-B”) The loss of function of RyR in the wing imaginal disc was used as a control (n=5) for the comparisons. (A-A”) Representation of immunohistochemistry data using standard deviation z plane projections along with cross-sectional views along a line parallel to the D-V boundary.(C-D”) Loss of function of Rho1 in the wing imaginal disc was introduced using commercially available Rho1^RNAi^ line (BL# 27727). (C-C”) Data representations similar to A-A”. (E) Rho1^RNAi^ (n=8) in the wing imaginal discs leads to an increased columnar cell height as compared to control discs (n=5). (F) Inhibition of Rho1 in the wing imaginal discs leads to flattening of the discs quantified through Menger curvature. A sample size of 5 and 8 was used for the control and Rho1^RNAi^ discs. (p-values of a student’s t-test included in plots) (G-I”) Using MS-1096 as a GAL4 driver allows for the knockdown to be existent in the dorsal compartment of wing imaginal disc. (G-H”’) Representation of immunohistochemistry data using standard deviation z plane projections along with cross-sectional views along a line parallel to the D-V boundary. (H–I”’) Cross-sectional views of the wing disc along lines parallel to the D-V boundary and located above and below the DV boundary as indicated by the yellow +δ and -δ lines in G.

We further investigated the impact on cell architecture of higher inhibition of Rho1 in the dorsal compartment using the ventral compartment as an internal control (Fig. 6G-I’’’). Cross-sectional views of the wing disc were examined that were parallel to the D-V boundary and located in the dorsal compartment as indicated by the yellow lines (+δ or - δ). Interestingly, we found that knockdown of Rho1 resulted in more basally located nuclei and a reduction of F-actin in the basolateral regions of cells. Surprisingly we also saw an increase in phosphorylated myosin II (pMyo-II) that is apically and basally localized and an increase in cell height. This could explain the decoupling of cell height and tissue curvature regulation. The implications for the deviations between experiments and simulations are explored in the discussion section.

### Tensile forces within the extracellular matrix maintain tissue bending

In the previous sections, we tested whether the passive tension of the ECM or actomyosin contractility beneath the nuclei were sufficient to bend the *Drosophila* wing disc. To further understand the mechanism underlying the bending shape, we also studied the tissue shape under perturbation conditions in both experiments and simulations (Fig. 7-9). Surprisingly, very little morphological change was observed when wing discs were incubated with either the Y-27632 ROCK inhibitor (not shown) or Latrunculin A (Fig 7A-7B). On face value, this result seemingly contradicts the inferred result of simulations presented in Fig. 5B. We thus investigated whether higher resistance of the ECMc to the applied forces can explain this result based on short-term pharmacological perturbations.

**Figure 7.**
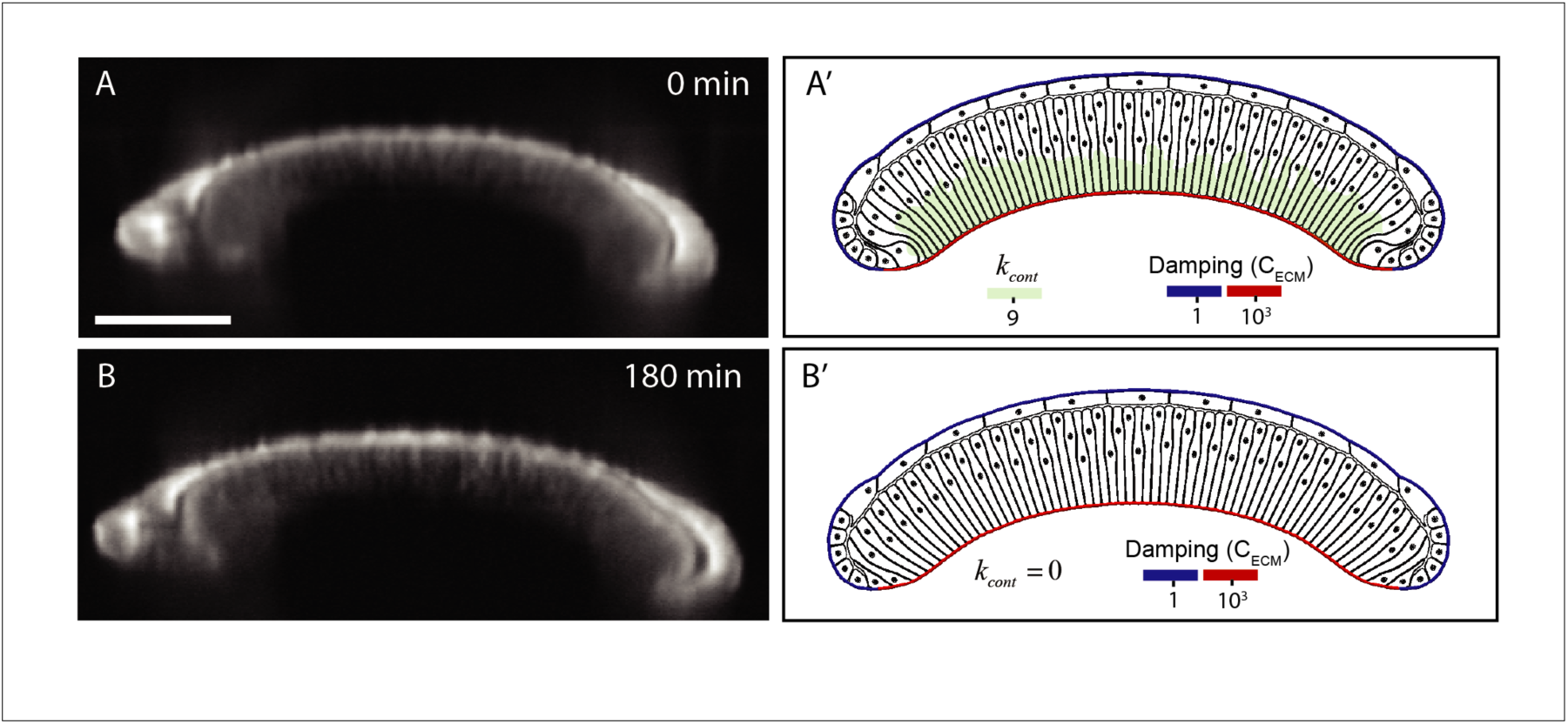
Comparison of experimental and simulated profiles demonstrates that ECM is sufficient to maintain profile of the tissue. A) Experimental profile before adding Latrunculin A to inhibit actomyosin. Scale bar is 50 μ*m*. The wing disc was stained with CellMask. Note that the imaging conditions required for live-imaging do not provide as fine resolution as for fixed images. A’) Computational model after the bent profile of wing disc is formed with higher levels of basal actomyosin contractility. B) Experimental profile three hours after addition of 4 µM Latrunculin to inhibit actomyosin. B’) Computational profile of wing disc showing that bend profile of wing disc is preserved even after removal of basal actomyosin contraction.

**Figure 9.**
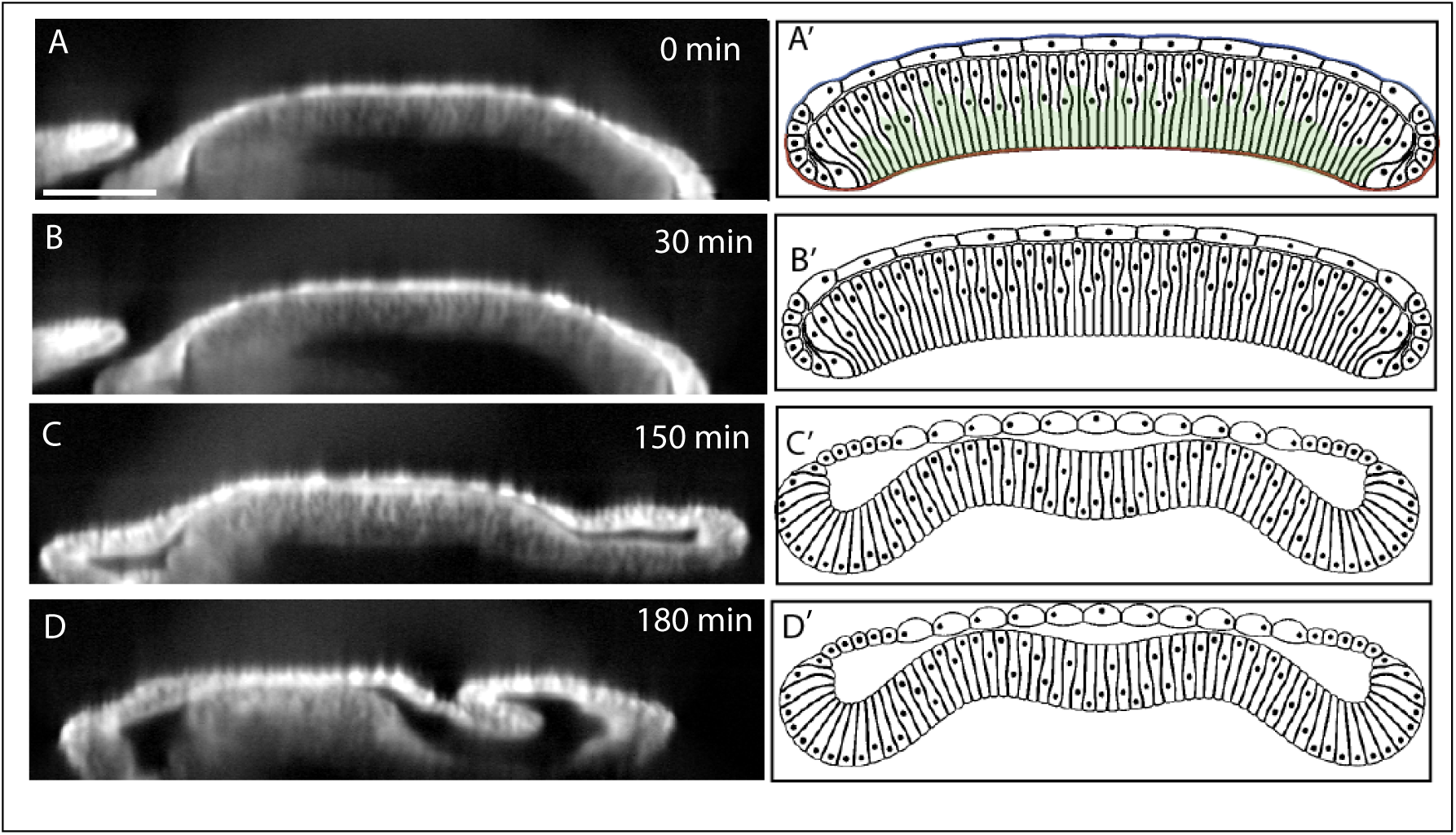
Enzymatic degradation of Collagen IV and actomyosin contractility in 96 h wing discs A-D) Enzymatic degradation of the wing disc in the presence of ROCK inhibitor to inhibit actomyosin contractility results in loss of adhesion between the squamous cells (top layer) and the central columnar cells. A’-D’) Computational simulation of acute removal of ECM, removal of adhesion between cell layers, and removal of actomyosin contractility components below the nuclear qualitatively match the experimental results. Experimental wing disc is stained with CellMask. Scale bar is 50 μ*m*.

Very recent experimental results support a significant role of the ECM in maintaining the shape of the tissue. Keller et al. demonstrated using finite element modeling and stretching experiments that the ECM is significantly stiffer than the columnar wing disc cells^39^. This provides additional evidence for compression by the ECM playing a defining role in maintaining the overall organ shape. In addition, Tozluoğlu et al. showed that ECM plays a major role in maintaining the folds in the wing disc in the orthogonal dorsal-ventral direction^39^.

Therefore, to explain the experimental observations, we hypothesized that after the bent profile of wing disc is “generated” by basal actomyosin contractility, the ECM becomes resistant to deformation. Thus, we infer that the relative passive prestrain that builds up in the ECM serves as a “memory” of tissue shape to preserve the bended shape of the tissue but does not actually generate the tissue shape. To represent this, we first ran a simulation to form the bent profile of wing disc and then computationally turned off the basal contraction from the simulation scenario while increasing the damping coefficient of the columnar-associated ECM (ECMc) by three orders of magnitude (Fig. 6A, B). This allows the ECM to preserve its shape and hence preserve the shape of the wing disc.

In comparison, we experimentally tested whether inhibition of the actin polymerization with Latrunculin A would impact the tissue shape in ex vivo organ cultures^15^. Surprisingly, we found that inhibition of actin polymerization had a minimal impact on tissue shape maintenance (Fig. 7A,B). This agrees with computational simulations indicating that a high mechanical resistance of the ECMc is sufficient to preserve the generated shape (Fig. 7A’, B’).

As further validation, we imaged the shape changes that occur when the ECM is chemically digested with collagenase in a 96 AEL wing disc. This perturbation resulted in inversion of the bent wing disc pouch with the margin area curving toward the squamous cells (Fig. 8C-D and Fig. 8C’-D’, SI Movies 1, 2). This confirms that the ECM is essential for maintaining the overall curved cross-sectional profile along the AP axis. Thus, the ECM serves as a ‘morphogenic memory system’. Additionally, these experiments provide insight into a third mechanism that contributes to the mechanical stability of the wing disc and the formation of the final shape. Without the ECM, the pulling force from the squamous cells to the columnar cells transmitted though marginal cells bends the entire tissue in the opposite direction of the final bending shape. The same mechanism results in an inverted bending profile near the marginal cells in Fig. 4A’.

**Figure 8.**
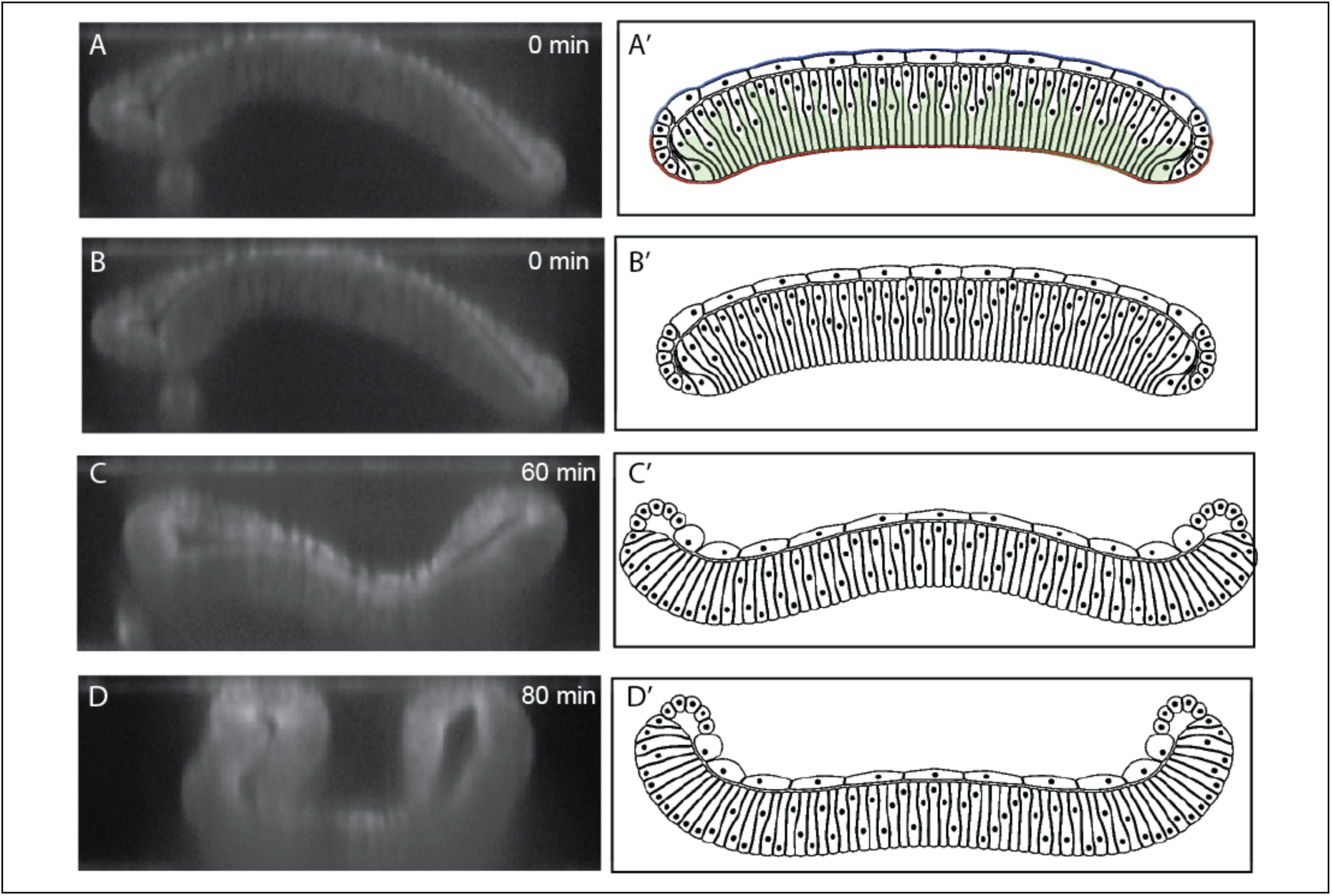
Enzymatic degradation of Collagen IV in 96 h wing discs. A-D) Enzymatic degradation of the wing disc results substantial difference in curvature profile of wing disc. A’-D’) Computational simulation of acute removal of ECM and removal of actomyosin contractility components below the nuclear qualitatively match the experimental results. Wing disc was stained with CellMask for experimental imaging.

The squamous cells become thinner as larval development proceeds. Hence, the pulling force from squamous cells, working against the desired bending profile, could be decreasing to allow the wing disc to position itself in a curved profile to facilitate the next steps of pupal morphogenesis. Therefore, this combined perturbation study provides evidence that the ECM is contributing to the bended shape of tissue by mechanically preserving, but not generating, the generated shape of the tissue.

### Dual ECM digestion/ROCK inhibition reveals significant apical adhesion between columnar and squamous cells

We used a ROCK inhibitor (Y-27632) at 1 mM concentration to deactivate actomyosin contractility in addition to the removal of ECM with collagenase to further investigate the relative contributions of actomyosin and the passive prestrain of the ECM (Fig. 9A-D and SI Video 2)^42, 43^. Experimental results revealed that the tight connection between the apical side of columnar cells near the dorsal-ventral compartment boundary and squamous cells in the pouch region is lost after inhibiting the actomyosin contractility in conjunction with collagenase treatment. A clear gap was generated between the two layers at the center, similar to genetic inhibition of Rho1 (Fig. 6). Similar gaps were also observed in simulation results (Fig. 9A’-D’) when adhesion between columnar and squamous cells, is reduced to zero in combination to deactivating the parameter representing basal actomyosin contractility and ECM. This suggests that actomyosin contractility not only pushes the nucleus and cytoplasm toward the apical surface in the columnar cells, but it is also involved in supporting the adhesion between columnar cells and squamous cells that is observed in localized regions. The adhesion between columnar and squamous cells are provided by microtubules extended between these two cell types^19^. Since microtubules are connected to the actin inside the cells, inhibition of ROCK may loosen the actin-microtubule network connection^44^. Hence columnar and squamous cells are not connected anymore, and a gap is formed between at the apical side between two layers. To summarize, this dual perturbation reveals a hidden mechanical role of the adhesion between the two layers of cells in the shape formation, and the relation between actomyosin contractility and adhesion at the apical side of columnar and squamous cells.

## Discussion

The regulation and maintenance of an organ’s shape is a major outstanding question in developmental biology. The *Drosophila* wing imaginal disc serves as a powerful system for elucidating design principles of the shape formation in epithelial morphogenesis^6, 7^. Yet, even simple epithelial systems such as wing disc are extremely complex. A tissue’s shape emerges from the integration of many biochemical and biophysical interactions between proteins, subcellular components, and cell-cell and cell-ECM interactions. How cellular mechanical properties affect tissue size and patterning of cell identities on the apical surface of the wing disc pouch have been intensively investigated^45, 46^. However, less effort has focused on studying the mechanisms governing the shape of the wing disc in the cross section. Both the significance and difficulty of such studies are due in part to the need to consider the composite nature of the material consisting of multiple cell layers and cell-ECM interactions as well as the elongated shape of columnar cells (Fig. 1B).

In this study, we iterated between experiments and newly developed computational model simulations to reverse-engineer the curved profile of the larval wing imaginal disc. This effort is aligned with an overall goal to elucidate general principles of morphogenesis. Namely, we developed a combined experimental imaging and biologically calibrated computational modeling approach to decouple the roles of actomyosin contractility, extracellular matrix tensile stress and cell-cell interactions involved in regulating organ morphology and nuclear positioning within cells. This resulted in the detailed quantification of the distribution of nuclei along the apical-basal axis, the height of the pouch, and the curvature of the pouch along the anterior-posterior axis.

Overall, our study defined the balance of forces determining the shape of the wing imaginal disc. The central insight obtained in this work suggests that actomyosin contractility is needed to generate the curved tissue profile along the anterior-posterior axis while tension within the ECM is sufficient and necessary for preserving the bent shape even in the absence of continued actomyosin contractility once the shape is generated. Accumulation of high levels of tensile stresses within the ECM along the basal side of columnar cells provides only a small positive contribution to generating the “dome-shaped” profile of the late 3^rd^ instar wing disc, and it cannot explain the asymmetric nuclei distribution of columnar cells. Therefore, passive tension built up in the ECM (perhaps that is generated over time due to differential growth patterns) does not contribute significantly to shape generation, at least for the bent profile of the disc cross-section along the AP axis. Hence, our study suggest another possible mechanism which is different than differential growth hypothesis^39^ to drive fold formation along the orthogonal dorsal-ventral axis. Perturbation studies that chemically dissolve collagen revealed that the ECM is essential to maintain the shape of the wing disc. Therefore, it provides “shape memory” for the tissue. Comparing experimental and computational results obtained under multiple perturbations demonstrated that actomyosin contractility was effective in generation of bended profile of wing disc, while ECM can effectively preserve the shape of the tissue.

Our results complement a recent report on the importance of lateral and basal contractility in the wing disc in the formation of several folds along the dorsal-ventral axis^15^. In contrast, our study focused on the bending along the orthogonal anterior-posterior axis. Further, our work also demonstrates a key mechanical role of asymmetrical nuclear positioning in defining the bent profile of the wing disc. Previous work has indicated that actomyosin contractility was important for the motion of mitotic cells within the wing disc^32^. Our results implicate basal actomyosin contractility in biasing all nuclei of the wing disc toward the apical surface. As the nucleus is stiffer than the cytoplasm, the positioning of nuclei near the apical surface may contribute to the formation of the shape of the disc.

Previous studies found that the peripodial membrane, a squamous epithelium that sits on top of the wing disc pouch, is connected to the pouch through microtubule-rich cellular extensions^19^. However, the implications of this interlayer connection are still not fully understood. Here we found that adhesion between the two layers keep the pouch and peripodial membrane together even under extreme morphological perturbations such as chemical digestion of the extracellular matrix. This provides general insight into latent interactions that ensure tissue integrity. Actomyosin contractility is needed to be maintained in this case for the two layers to remain adherent in localized regions such as the dorsal-ventral compartment boundary where the distance between the two layers is minimal. Separation of these two layers occurred with dual perturbations of collagenase and ROCK inhibition. This is likely due to the need of basally directed actomyosin contractility to keep pouch cells in close apposition to the peripodial membrane. Thus, adhesion between the apical surfaces of intra-organ layers depends on the maintenance of ROCK-driven actomyosin activity in the tissue to prevent delamination. Consequently, the spatiotemporal patterning of actomyosin contractility provides additional, indirect roles of actomyosin activity in maintaining tissue integrity that become apparent only through a series of perturbation experiments. From the collagenase-treatment experiment, we also uncovered the pulling force from squamous cells over the columnar cells through marginal cells as a third mechanism that acts in an antagonistic way to the ECM tensile stress and actomyosin contractility. This pulling force results in the wing disc inversion along the AP cross-section of the tissue observed when the ECM is removed by collagenase.

Genetic inhibition of Rho1, a key regulator of actomyosin contractility, confirmed the role of this pathway in defining tissue curvature. Analysis further showed a qualitative decrease in F-actin levels in basolateral regions of the columnar cells. However, as previously noted ^13^, the height of the cells is actually increased when Rho1 activity is increased. We found high levels of activated myosin near the basal and apical surfaces of columnar cells showing this height increase compared to controls (Fig. 6). This suggests that other regulators of Myosin II are involved in specifying the asymmetry in actomyosin contractility. Local contractility at the apical and basal surface may play important roles in cell height control, which computational simulations could help to address in the future. Other Rho family GTPases such as Cdc42 have been implicated in cell height regulation^47^, and their roles in defining subcellular contractility and overall tissue shape require further elucidation.

Larval wing disc development serves to generate the precursor to the fully formed wing. Wing morphogenesis occurs during subsequent pupal development and requires significant remodeling of the extracellular matrix^48, 49^. Therefore, the mechanisms tested in this paper explain the emergence of the key initial input geometry of the wing disc leading up to eversion and wing morphogenesis occurring during pupal development. Lastly, a combination of the multi-scale subcellular element modeling environment described in this paper and specially designed experiments can be readily extended to generate and test hypothesized novel mechanisms of the wing disc eversion process and other epithelial systems, including organoids, that consist of several cellular and ECM layers.

## Materials and Methods

### Fly culture and developmental staging

Details on fly husbandry and developmental staging, wing disc immunohistochemistry and imaging are provided in SI methods.

### Tissue surface and nuclei segmentation

In-house Matlab scripts were written to facilitate tissue orientation and segmentation. This program allows the user to trace the long axis and provides the corresponding cross section on which the user segments the apical and basal surfaces. Nuclei within these cross-sections were automatically segmented (SI Fig. 2). Once surface and nuclei segmentations were performed, the program calculates the local curvature of the basal surface as described in Fig. 2F as well as the fractional nuclei positions (Fig. 2B).

### Computational model

A novel multi-scale subcellular element (SCE) model was developed and calibrated using experimental data to simulate the cross-sectional profile of the wing along the anterior-posterior axis. The model includes three types of nodes to represent mechanical properties of the cell nucleus, membrane and ECM, with interactions between nodes prescribed by different energy functions. The energy functions and related biological justifications are given in Table 1 and shown in Fig. 10A. Model describes columnar cells (Fig. 10A), boundary cells (Fig. 10B) and squamous cells (Fig. 10C). ECM nodes, which surround basal sides of epithelial cells, are subdivided into three types. ECM nodes connected with columnar cells, connected with boundary cells and boundary cells are called ECMc, ECMbc and ECMs, respectively. In this paper, we tested multiple mechanistic scenarios prescribing different mechanical properties to different regions of ECM and different levels of basal actomyosin contractility.

**Table 1:** Potential energy functions used in the developed subcellular element model.

| Energy term | Type | Physical representation |
| --- | --- | --- |
| $E_v$ | Morse | Volume exclusion |
| $E_{nuc}$ | Morse | Size of nucleus |
| $E_{adhL}$ | Spring | E-cadherin (cell-cell adhesion) |
| $E_{adhB}$ | Spring | Integrin (cell-ECM adhesion) |
| $E_{adhA}$ | Spring | Adhesion between columnar and squamous cells |
| $E_{cont}$ | Spring | Actomyosin contractility inside the cells and beneath the nucleus |
| $E_{memb}$ | Spring | Actomyosin contractility at the cell cortex and membrane stiffness |
| $E_{ecm}$ | Spring | Extracellular matrix stiffness |
| $E_{vol}$ | Lagrange multiplier | Volume conservation of cytoplasm for each cell |

The positions of all nodes describing different sections of the ECM and different types of cells are updated by using different potential energy functions listed in Table 1 and based on Langevin equations.

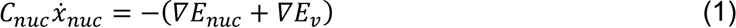

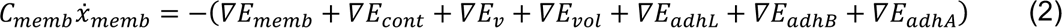

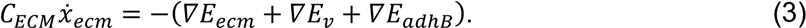

The Langevin equation assumes that cell motion occurs in in the overdamped regime^50^. Cell-cell adhesion and cell-ECM adhesion model parameters were calibrated using experimentally obtained values of adhesion for epithelial cells^51^ since it was reported that the level of cell-cell adhesion for epithelial cells is in the same range as the cell-ECM adhesion^52^. Size and number of cells in the model were calibrated using experimentally observed shapes of epithelial cells (see SI Table 1). The ranges of all model parameters are provided in SI. 7.

## Acknowledgements

ML and JZ were supported in part by the NIH Grant R35GM124935 and NSF Awards CBET-1553826. MA, WC and AN were partially supported by the NSF Grant DMS-1762063 through the joint NSF DMS/NIH NIGMS Initiative to Support Research at the Interface of the Biological and Mathematical Sciences. MA was partially supported by the NIH Grant UO1 HL116330. The authors acknowledge support from the Notre Dame Integrated Imaging Facility and CRC Computing center. We would like to thank members of the Dr. Zartman’s and Dr. Alber’s groups for their helpful feedback. We thank Jamison Jangula for significant technical assistance during the initial project stages and Andrew Whitaker for help with post processing of computational results.

