## Supplementary material for "Epithelial organ shape is generated by patterned actomyosin contractility and maintained by the extracellular matrix": SI Text

### SI Methods

**SI.1 Additional fly culture details.** Flies were raised at 25 °C and a twelve-hour light cycle unless specified otherwise. For all staging experiments, adult flies were used three days after pupal eclosion. These adults were placed in cages containing grape agar plates and yeast paste for four hours. Eggs from this collection period were allowed to mature on the agar plates for 24 hours after which first instar larva were transferred to new agar plates. The time used for the developmental age was determined from the midpoint of the egg collection period. Fly stocks include: The *Drosophila melanogaster* line Oregon-R was used in all staging experiments (SI Fig.1). Vkg:GFP expressing flies were used for imaging of the ECM. MS1096-Gal4 were used to generate results in Figure 6.

Live images were collected from discs cultured in Grace's medium (ThermoFisher, 11595030) with low ecdysone (Sigma, H5142) [1]. Live imaging experiments were performed for up to four hours with 100-200, 1 $\mu$m slices taken across the z-direction every 5 minutes. Note that the imaging conditions required for live-imaging do not provide as fine resolution as for fixed images. 1 mM Y-27632 (Selleck Chemicals LLC, S1049) was used for ROCK inhibitor experiments [2,3]. 3 mg/ml Collagenase (Worthington Biochemical, LS004194) was used for ECM inhibition experiments. 4 μM Latrunculin A (Sigma-Aldrich, L5163) was used to inhibit actin during live imaging[4]. Finally, 1 μl/ml of Cell Mask (ThermoFisher, C10046) was used to visualize the shape of the disc during treatment.

**SI.2 Wing disc immunohistochemistry and mounting.** Wing imaginal discs from 3rd instar *Drosophila* larva were dissected in intervals of 20 min. Dissected wing discs were then fixed in ice cold 10% neutral-buffered formalin (NBF) for 20 min in PCR tubes. Immediately following fixation, wing discs were rinsed three times with fresh PBT (PBS with 0.03% v/v Triton X-100). Tubes containing wing discs were then placed on a nutator for 10 minutes at room temperature and then rinsed again with PBT; this step was repeated for a total of three nutation/rinsing intervals. PBT from the final rinse was removed and 200 µL of 5% normal goat serum (NGS) in PBS was added to each PCR tube. Tubes were then placed on a nutator for 30 minutes at room temperature. NGS was then removed and 200 µL of a primary antibody mixture, prepared in 5% NGS solution, was added. Tubes were then placed on a nutator at 4˚C overnight. Three quick rinses were then performed as was done after fixation. Tubes were placed on a nutator for 15 minutes followed by PBT rinsing; repeated for a total of three nutation/rinsing intervals. PBT from the final rinse was then removed and 200 µL of a secondary antibody mixture, prepared in 5% NGS solution, was added. Tubes were then placed on a nutator for 2 hours at room temperature. Three quick rinses were then performed as before. Tubes were then placed on a nutator for 20 minutes at room temperature and then rinsed with PBT for a total of three intervals. Wing discs were then either kept in tubes at 4˚C or mounted immediately.

**SI.3 Mounting of stained wing discs**. Wing discs were mounted between 24 x 60 mm and 22 x 22 mm glass cover slips with VECTASHIELD® (Vector Laboratories). For images in which the natural curvature of the wing disc was desired, Scotch tape was used as a spacer between the two cover slips. Clear nail polish was used to seal in VECTASHIELD® and to adhere cover slips together.

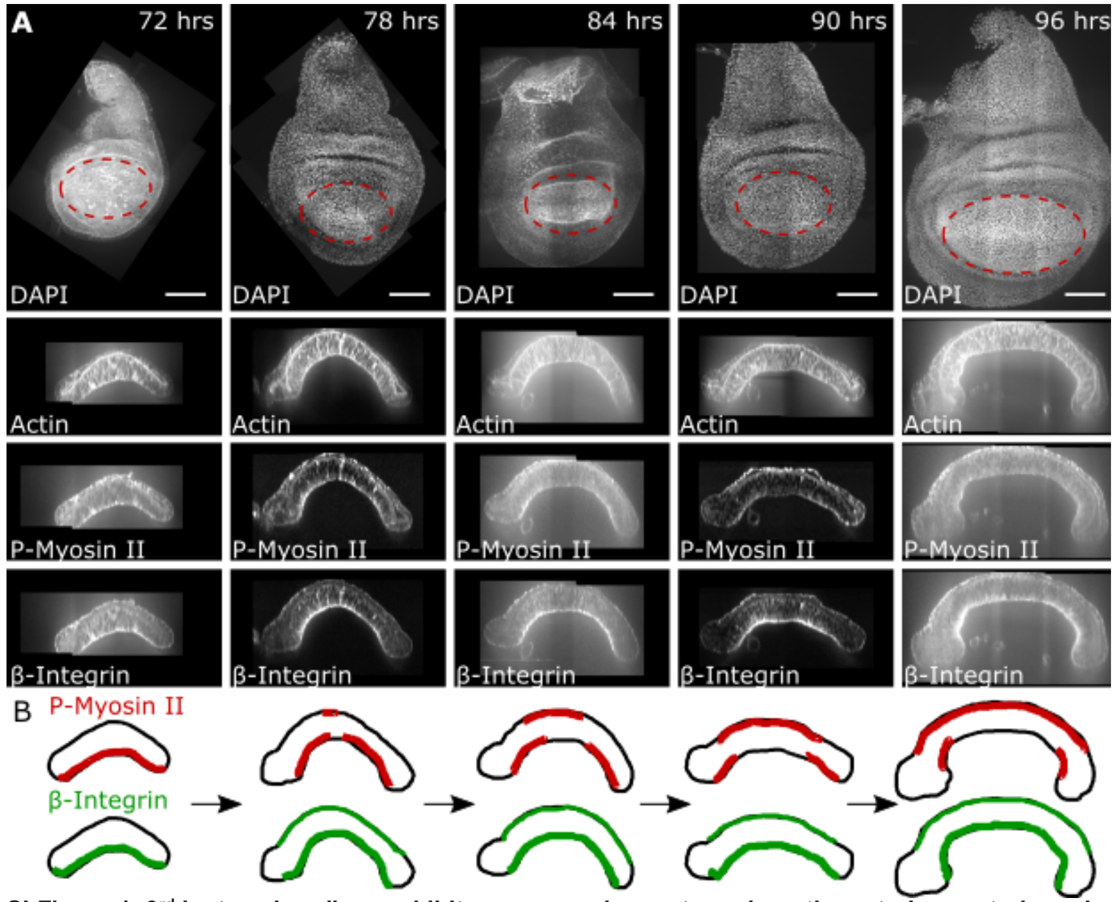


**SI Figure 1. 3rd instar wing discs exhibit pronounced curvature along the anterior-posterior axis. A)** Wing disc development from 72 to 96 hours AEL. The first row shows z-projections of nuclear signals (DAPI). Rows 2-4 are cross-sections along the anterior-posterior axis showing F-actin (phalloidin), P-Myosin II, and $\beta - \mathrm{integrin}$. Scale bars are 50 $\mu m$. **B)** Schematic showing asymmetric of P-Myosin II and β-Integrin for each timepoint in **A**. The genotype for SI Figure 1 is Ore R.

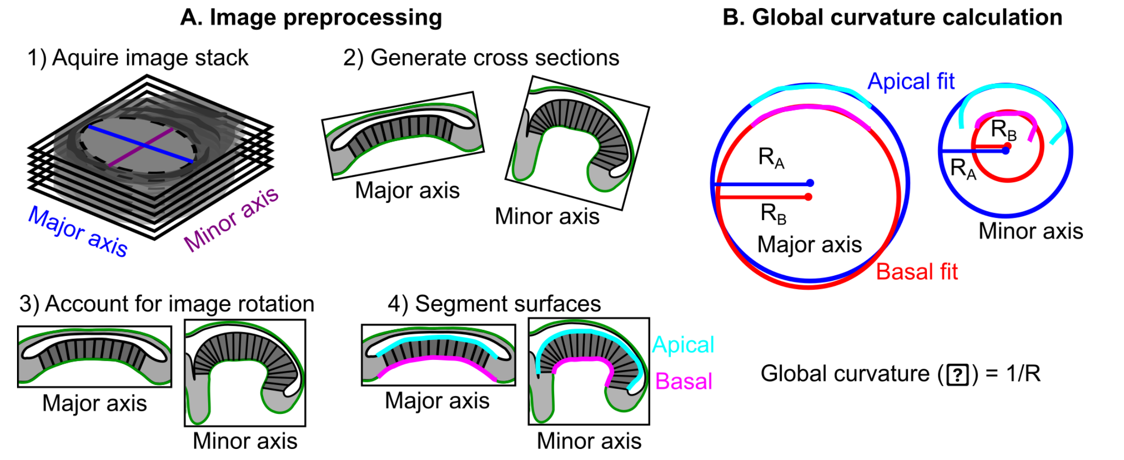


**SI Figure 2. Description of the pipeline developed to extract the cross-section of wing discs and quantification of global curvature as the metric to compare different phenotypes. A)** Pipeline to extract apical and basal surfaces along the major and minor pouch axes. **B)** Global curvature is obtained by fitting an arch of a circle through a set of data points selected from the surfaces.

**SI. 4 Additional image processing methods**. Proper orientation is critical for determining tissue thickness, since misalignment in one plane affects measurements taken from images of orthogonal planes. To prevent measurement errors associated with misalignment, we developed a pipeline for quick image orientation and surface segmentation (SI Fig. 2). This pipeline was used to extract apical and basal surfaces along the major and minor pouch axes. We chose global curvature as a simple metric to quantify curvature development during wing disc growth. Global curvature is calculated by fitting an arch of a circle through a set of data points and taking the inverse of the circle's radius. Global curvatures were calculated for the segmented apical and basal surfaces (SI Fig. 2).

To quantify the nuclear position in the cross-section of wing discs, we developed a pipeline to segment nuclei from images of DAPI (DNA marker) stained tissues (SI Fig. 3). This pipeline implements various image processing steps followed by calculation of the centroid of each nucleus. The statistics obtained by this quantification are shown in SI Fig. 3B.

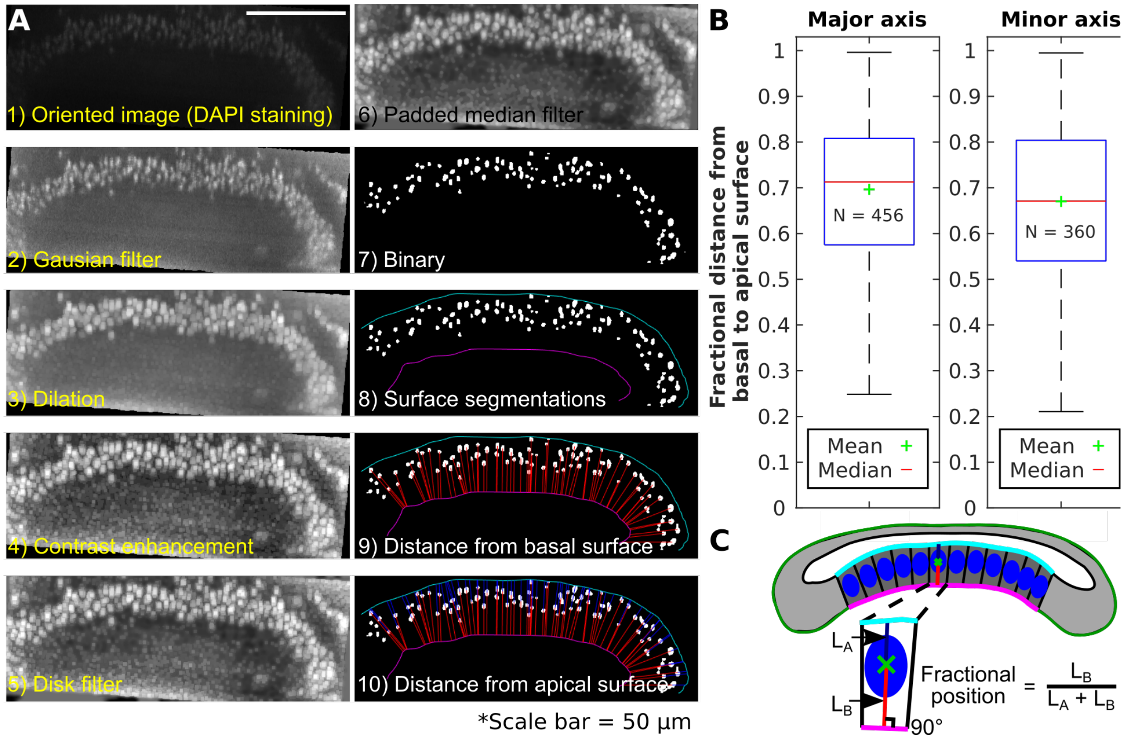


**SI Figure 3. Quantification of apical nuclear bias. A)** The pipeline implements various image processing steps (1-6) followed by extracting the centroid of each nucleus (7). Lines perpendicular to the basal surface, obtained in (8), are then generated starting from each nuclear centroid (9). The distances from each centroid to the apical and basal surfaces are then calculated as the fractional positions of nuclei (1). **B)** Boxplots of the fractional distances of nuclei from basal to apical surfaces. **C)** Definition of fractional position.

To quantify the difference in tissue's thickness due to ECM degradation, distances were extracted from the apical to basal surface through each segmented nuclei Collagenase treatment experiments were performed to decouple the mechanical contribution of the extracellular matrix from all other tissue components. Collagenases are enzymes that break the peptide bonds in collagen, a structural protein of the ECM. Viking (Vkg) is a subunit of collagen IV, and we expressed Vkg::GFP in the wing disc to visualize collagen within the epithelium. We incubated wing discs in live culture media containing 3.4 mg/ml collagenase for 1 hour. Before media preparation, collagenase was dissolved in Phosphate-buffered saline (PBS). To ensure that differences between control and treated wing discs were due to collagenase, we added an equivalent volume of PBS to the control media. We found that thickness differences were significant between the untreated and collagenase treated conditions (SI Fig. 4).

SI Figure 5 shows the distribution of phosphorylated Myosin II in cross-section (A), as well as in individual planes encompassing the squamous peripodial cells (B), the columnar pouch cells near the apical surface (C) and near the basal surface of the columnar pouch cells (D).

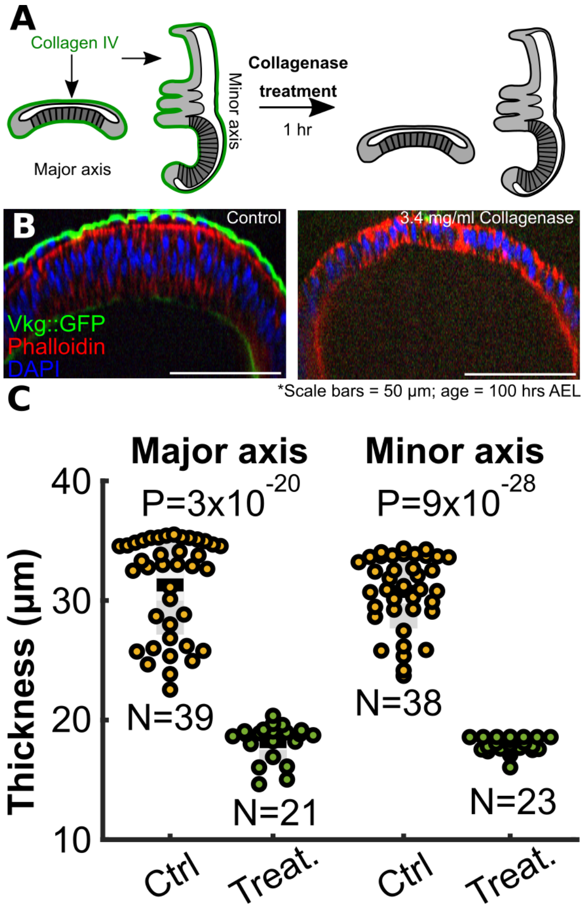


**SI Figure 4. Chemically based ECM removal**. ECM degradation by collagenase treatment (A) resulted in a visually distinct decrease in thickness of columnar epithelium (B) as quantified in (C).

To calculate the Menger curvature for the intensity correlations, a region of interest defining the wing imaginal disc was manually selected using an open source MATLAB package ‘roispline’ [5]. A curve defining the apical surface of wing disc was then separated from the wing disc boundary points defined by the fitted spline. An equally spaced fixed number of boundary points were then selected from the curve as indicated by a red dot in Fig 2G. For any set of $i^{th}$ points, we defined Menger curvature as the reciprocal of the radius of the circle passing through the three points [6]. Fluorescence intensity around any $i^{th}$ point was defined as the averaged integrin intensity along the curve defined between the $i\text{-}1^{th}$ and $i{+}1^{th}$ point. The circle is fit using an inbuilt MATLAB function ‘circumcenter’.

Then to calculate local curvature points of the wing disc boundary were selected interactively using custom built MATLAB functions. Between any two consecutive points, a natural cubic spline was fit following which the segment defining apical boundary was separated from the wing disc. The whole methodology is similar to one described above. We then used the spline $S_i$ defined between points $X_i$ and $X_{i+1}$ to interpolate sufficient number of points for estimation of derivatives and double derivatives which are further used for curvature calculation.

.

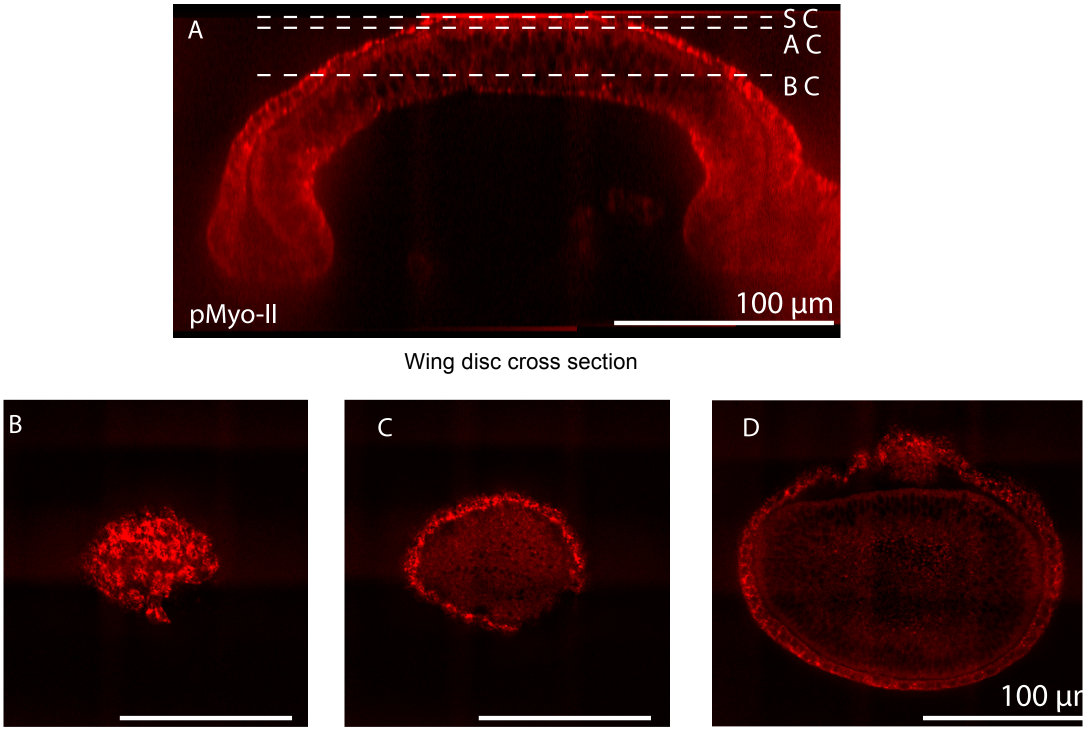


Wing disc cross section

Squamous Cells (SC) Apical Columnar (AC) Basal Columnar (BC)

**SI Figure 5.** Distribution of phosphorylated Myosin-II (pMyo-II) in cross-section and in individual z-slices of wing discs. **A)** pMyo-II distribution in the cross-section of the wing disc along the anterior-posterior axis, **B)** a single z-slice covering the squamous peripodial cells. **C)** pMyo-II distribution near the apical surface of the pouch cells, **D)** the basal surface of pouch cells.

**SI.5 Computational model description**. In this paper, a novel subcellular element model (SCE) is introduced to decouple direct and indirect effects of actomyosin-generated forces, nuclear positioning, passive tension in the extracellular matrix (ECM), and adhesion in shaping *Drosophila* wing imaginal discs. This SCE model includes two sets of nodes. The first set of nodes includes intra-cellular structural components such as membrane nodes and nucleus nodes. While, the second set of nodes represents extra-cellular structural components such as ECM nodes. Different potential energy functions are used to model interactions between the same type or among different types of nodes (Fig. 10). More specifically, $E_v$ is the Morse potential energy function describing volume exclusion between different types of nodes. $E_{nuc}$ is another Morse type potential defining the size of the nucleus. $E_{adhL}$ is a pairwise spring potential describing the interaction between sufficiently close membrane nodes of neighboring cells along the lateral sides. $E_{adhB}$ represents the force provided by integrin, a transmembrane protein, as cells adhere to the ECM. $E_{adhA}$ represents the apical connection between columnar and squamous cells[7]. $E_{cont}$ represents actomyosin contraction beneath nuclei of columnar cells. $E_{cont}$ is modeled by a pairwise spring potential between lateral membrane nodes, yielding the apically biased nuclear positioning in columnar cells (SI Fig. 3). $E_{memb}$ represents cortical stiffness associated with lipid bilayer for each cell. $E_{ECM}$ represents ECM stiffness provided mainly by collagen. The Lagrange multiplier energy function $E_{vol}$ is used to guarantee the conservation of volumes of individual cells (mainly cytoplasm) as follows:

$$E_{vol} = k_{vol}(\Omega - \Omega_0)^2, \quad \text{(SI 1)}$$

where $\Omega$ is the current volume of the cell and $\Omega_0$ is the prescribed cellular volume. The coefficient $k_{vol}$ determines the strength of the enforcement of the volume constrain. Following previous studies[8,9], we choose a specific value $k_{vol}$ (SI Table 3) such that the cell volume remains conserved in all simulations with 90% accuracy.

Equations 1-3 in the main text of the paper describe motion of different types of nodes, which is determined by the descent gradient of the corresponding potential energy function. Since all potentials are continuous functions, those gradients can be derived analytically. For example, $E_v$ , a Morse energy potential function corresponding to volume exclusion, has the following form:

$$E_v = U_v exp\left(\frac{-|\boldsymbol{x}_i - \boldsymbol{x}_j|}{\zeta_v}\right) - W_v exp\left(\frac{-|\boldsymbol{x}_i - \boldsymbol{x}_j|}{\gamma_v}\right) \quad \text{(SI 2)}$$

Then the gradient of the $E_v$ can be calculated as follows:

$$\nabla E_v = \left(\frac{-U_v}{\zeta_v} exp\left(\frac{-|\boldsymbol{x}_i - \boldsymbol{x}_j|}{\zeta_v}\right) + \frac{W_v}{\gamma_v} exp\left(\frac{-|\boldsymbol{x}_i - \boldsymbol{x}_j|}{\gamma_v}\right)\right)\frac{\boldsymbol{x}_i - \boldsymbol{x}_j}{|\boldsymbol{x}_i - \boldsymbol{x}_j|}, \quad \text{(SI 3)}$$

where $U_v$ , $\zeta_v$ , $W_v$, and $\gamma_v$ are Morse coefficients, and $\boldsymbol{x}_i$ and $\boldsymbol{x}_j$ are vectors describing locations of nodes i and node j. All vectors describing location of nodes are started from zero. Knowing minus of gradient of potential energy is equal to force, equation SI 3 describes the impact of the force applied to node i by node j due to existence of the $E_v$ between these two nodes.

Similarly, all terms on the right-hand sides of the Equations 1-3 are calculated using potential energy functions listed in Table 1. Terms on the left-hand sides of the Equations 1-3 are discretized by using explicit Euler method and solved on a cluster of graphical processing units (GPUs) to accelerate the computation. For more detailed information about discretization and GPU implementation, please see [10].

By using the SCE model developed above, we investigated roles of ECM and basal actomyosin contractility in bending the wing disc tissue along the AP axis. Regarding the role of ECM, we compared the modeling results obtained with differential tensile stresses in the ECM which is modeled by assuming different values for $L_{0_{ECMc}}$ and $L_{0_{ECMs}}$ (see Fig. 10). Tensile forces in the ECMc and ECMs are defined as below:

$$F_{ECMc} = k_{ECM}\left(L - L_{0_{ECMc}}\right), \quad \text{(SI 4)}$$

$$F_{ECMs} = k_{ECM}\left(L - L_{0_{ECMs}}\right). \quad \text{(SI 5)}$$

Since $k_{ECM}$ is identical for ECMc and ECMs, lower value of $L_{0_{ECMc}}$ compared with $L_{0_{ECMs}}$ leads to higher initial tensile stress in ECMc compared with ECMs.

The role of actomyosin contractility was investigated by varying the stiffness of the springs connecting lateral nodes below nuclei of columnar cells in the model (Fig. 10). As the spring stiffness is increased, the tendency of columnar cells to squeeze beneath the nuclei is also enhanced based on the following equation

$$F_{cont} = k_{cont}\left(L_{cont} - L_{0_{cont}}\right). \quad \text{(SI 6)}$$

To identify each mechanism in bending the tissue, we separately varied the ratio between $F_{ECMc}$ and $F_{ECMs}$ and $F_{cont}$. The resulting tissue shape evaluated in terms of overall curvature, relative nuclear positions in individual cells and the height of columnar cells was compared with the experimental data as shown in Fig. 4 and Fig. 5.

To guarantee that simulation of the tissue shape reaches steady state, we checked convergence of the temporal profile of the curvature generated by the model. A flat sheet of cells was used as the initial condition with zero global curvature (Fig. 4A and Fig. 5A). The parameters for initial shape of the tissue is given based on measurements of different properties of cross section of wing disc in the current research and it is shown in SI Table 1.

**SI Table 1.** Calibration of the model using experimental data over 96 hours of development of the cross-sectional profile of the wing along the anterior-posterior axis.

| **Properties** | **Experiment (this work)** | **Simulation** |
|---|---|---|
| Number of columnar cells | 60-70 | 65 |
| Columnar cells width | 2-3 $\mu m$ | $2.5 \mu m$ |
| Columnar cells height | 25-35 $\mu m$ | 25 $\mu m$ |
| Number of squamous cells | 10-15 | 10 |
| Squamous cells width | 15-18 | 16 |
| Squamous cells height | 3-5 | 4 |
| Nucleus distribution of columnar cells | $60\% \pm 16\%$ | $70\% \pm 16\%$ |

The simulation was run for 120,000 time steps, which resulted in near convergence to a stable, final shape (SI Fig. 6). The asymptotic fitting regression was also plotted to quantify the final value of the global curvature. It can be seen that asymptotic fit regression model fits well with the simulation results, showing that reliable convergence is obtained.

The code for this computational framework was developed in CUDA C++ and it is available on github: https://github.com/AliNemat/EpiScale_CrossSection.git.

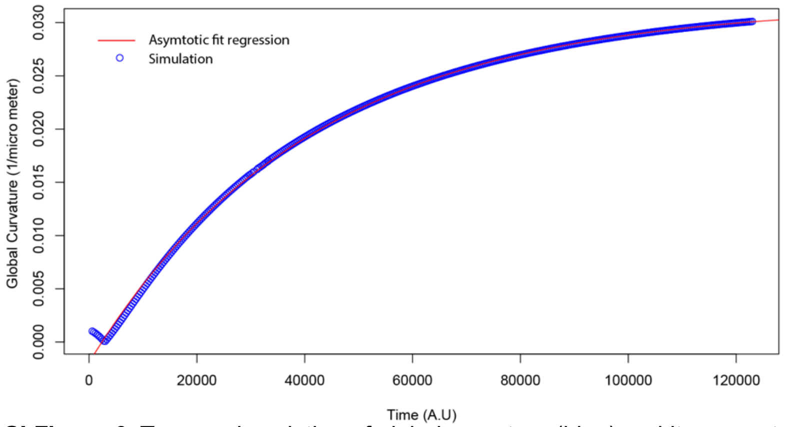


**SI Figure 6.** Temporal evolution of global curvature (blue) and its asymptotic fitting regression curve (red) for a simulation leading to the steady state shape in the wild-type simulation.

**SI.6 Latin hypercube sampling (LHS) method and sensitivity analysis.** The Latin hypercube sampling (LHS) method was applied to perform the sensitivity analysis of the bending shape of tissues with respect to multiple parameters in the computational model [11]. LHS method is one of the most efficient sampling methods for sensitivity analysis, especially when the number of parameters is large [12]. In particular, we chose average diameter of the cellular nuclei, the ratio between the tension of ECM connecting with columnar cells and squamous cells ($F_{ECMc}/F_{ECMs}$), and the level of actomyosin contractility below nuclei in columnar cells ($K_{cont}$) as the inputs for the sensitivity analysis. In the output, we compared the global curvature of the basal side of columnar cells, mean position of nuclei in columnar cells and the mean height of columnar cells. Minimum and maximum values for each of these three parameters are chosen using LHS method to obtain final temporal global curvature. In the LHS sensitivity analysis, the range of each parameter was divided into 10 bins and exactly 10 samples are chosen following the uniform random distribution such that there is exactly one sample selected from each bin for every parameter. A typical result of 10 sets of parameters obtained by this sampling method is shown in SI Fig. 7.

The code is available on github: https://github.com/AliNemat/LatinHyperCube.git.

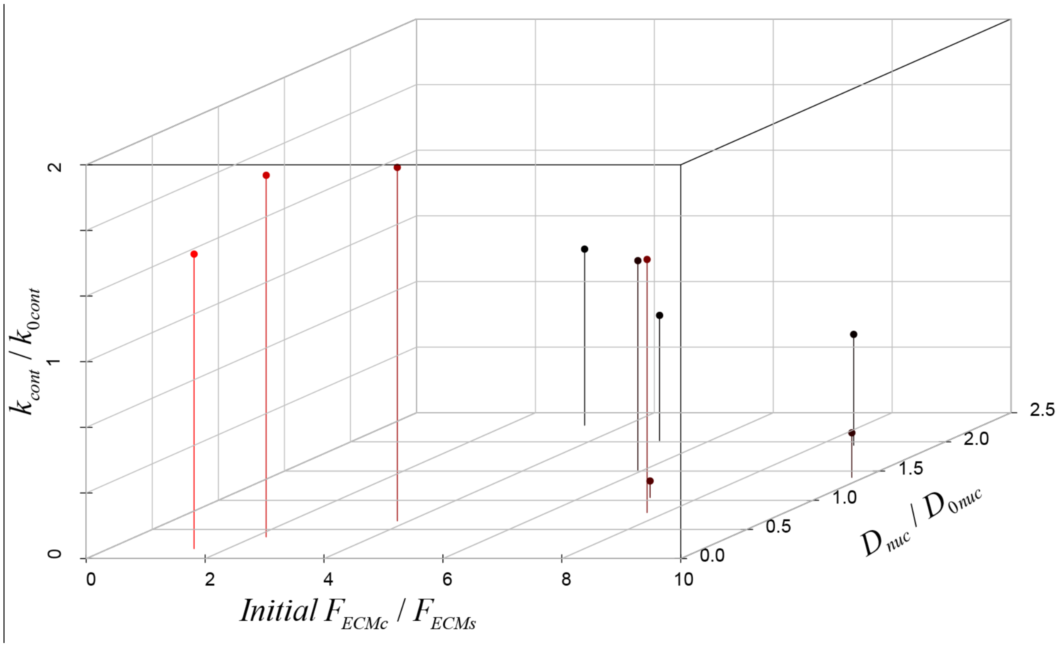


**SI Figure 7.** Hypercube method demonstrates sensitivity of the bending shape with respect to nucleus diameter, ratio of the tensions of ECM connected with the columnar cells and squamous cells as well as actomyosin contractility beneath nuclei.

After obtaining sampling parameter set using LHS method, we applied partial correlation coefficient method[11] to assess the relation between three input parameters and three output quantities measured from the computational model simulations[11]. In particular, the global curvature was calculated by using best circle fitting method which was also used in image analysis of experimental data (SI Fig. 2). The average height of columnar cells is calculated by the average distance between apical and basal nodes of columnar cells. Finally, the average nuclear position in columnar cells are calculated as below

$$\overline{loc}_{nuc}(\%) = \frac{1}{n}\sum_{i=1}^{n}\frac{|\boldsymbol{x}_{nuc}(i) - \boldsymbol{x}_{bas}(i)|}{|\boldsymbol{x}_{api}(i) - \boldsymbol{x}_{bas}(i)|}, \quad \text{(SI 7)}$$

where n is the number of columnar cells, $\boldsymbol{x}_{nuc}$ is the vector defining the position of the nuclear center of each cell, $\boldsymbol{x}_{bas}$ is the vector defining the basal location of each columnar cell and $\boldsymbol{x}_{api}$ is the vector defining the apical location of each columnar cell. In the partial correlation coefficient method, the relationship between an input parameter and an output measurement is characterized through removing the linear correlation between this output measurement and all other input parameters. The inputs and outputs for the sensitivity analysis are listed in SI Table 2.

**SI Table 2.** Inputs and outputs in the partial correlation coefficient study.

| Input parameters | Output parameters |
|---|---|
| $x_1 = x_{ten} = F_{ECMc}/F_{ECMs}$ | $y_1 = y_{curv}$ |
| $x_2 = x_{nucD} = D_{nuc}/D_{0nuc}$ | $y_2 = y_{nuc}$ |
| $x_3 = x_{cont} = k_{cont}/k_{0cont}$ | $y_3 = y_h$ |

Then the partial correlation coefficient between $x_i$and $y_j$ is the correlation coefficient between the residual of the input parameter, $x_i - \hat{x}_i$, and that of the output measurement, $y_j - \hat{y}_{ji}$, where

$$\hat{x}_i = c_0 + \sum_{\substack{p=1 \\ p\neq i}}^{3} c_p x_p, \tag{SI 8}$$

$$\hat{y}_{ji} = b_0 + \sum_{\substack{p=1 \\ p\neq i}}^{3} m_{jp} x_p. \tag{SI 9}$$

The overall result of the sensitivity analysis based on partial correlation coefficient method is shown in SI Fig. 8. Among all three input parameters, there is strong positive linear relationship between the actomyosin contractility and all three output measurements (SI Fig. 8C, C', C").

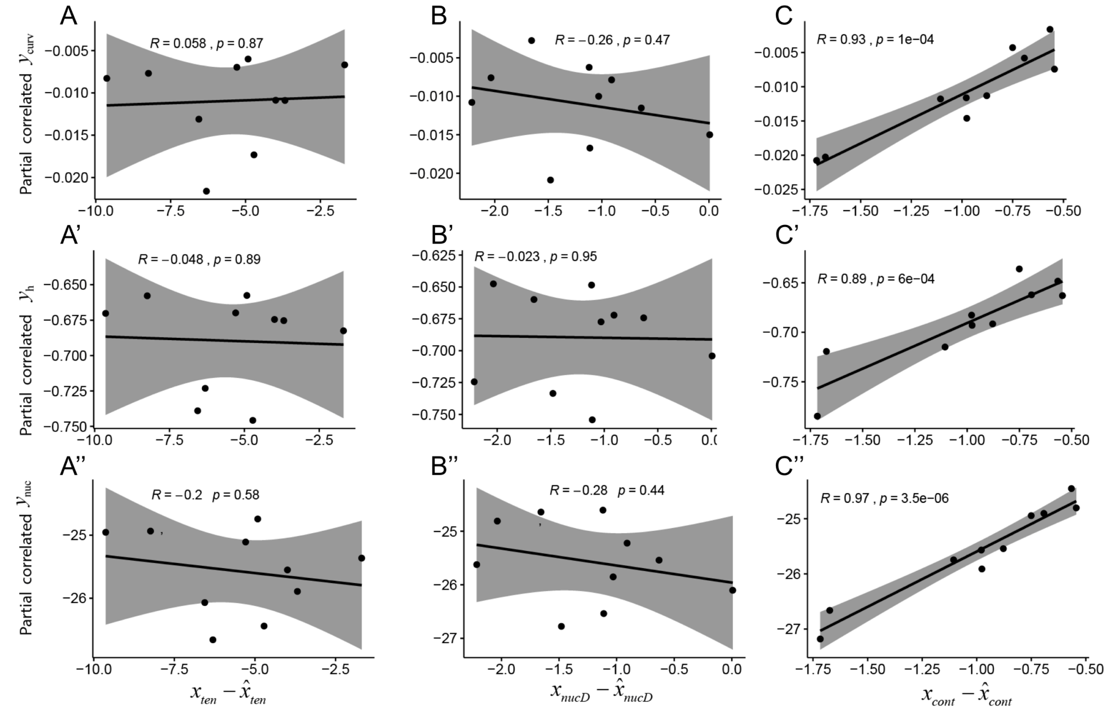


**SI Figure 8.** Correlation between the outputs: global curvature of the tissue (A, B, C), average nucleus position (A', B', C') and average height of columnar cells (A", B", C"), and inputs: ECM relative tension (A, A', A"), nucleus average diameter (B, B', B") and basal contraction of pouch cells (C, C', C") obtained by using partial correlation coefficient method.

In addition, nuclear diameter is weakly negatively correlated with the global curvature and cell height (SI Fig. 8B, B', B"), while the ECM tension has weak negative correlation with the cell height (SI Fig. 8A, A', A"). No linear correlation was observed for all other pairs between inputs and outputs. The corresponding quantification of the pairwise correlation is shown in SI Fig. 9. We can see that the correlation with actomyosin contractility is highest for all three outputs and the nuclear diameter is only correlated with the global curvature and columnar cell height negatively. The ECM tension has no significant correlation with all three outputs. These quantifications are all consistent with results shown in SI Fig. 8. Overall, the sensitivity analysis also confirmed that basal actomyosin contractility is enough to induce tissue bending while passive tension built up within the ECM is not enough to bend the tissue in the *Drosophila* wing disc along the anterior-posterior axis.

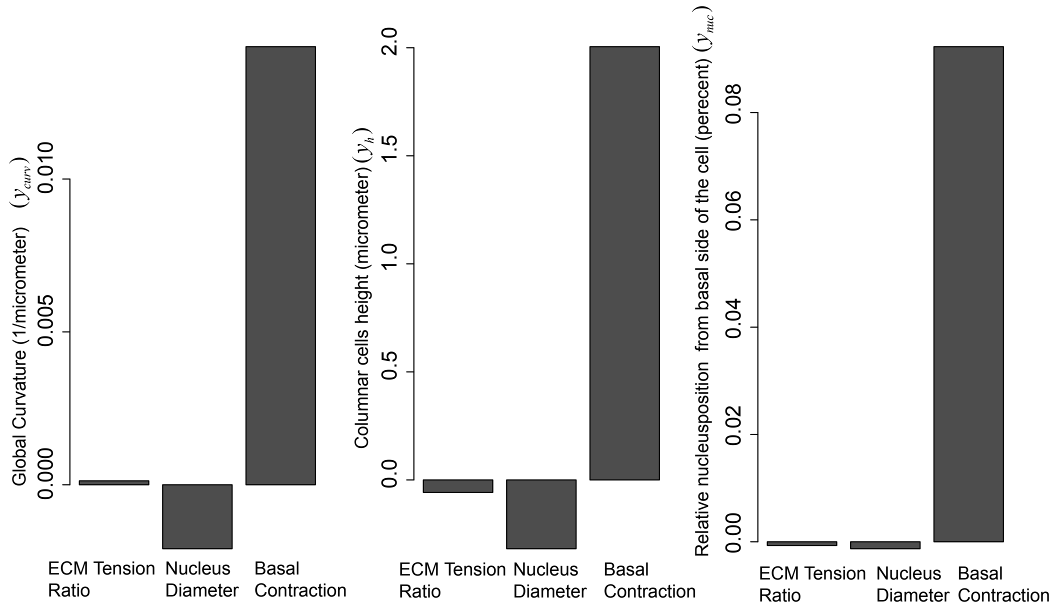


**SI Figure 9.** Sensitivity of global curvature of the tissue, average nuclear position and average height of columnar cells with respect to changes in ECM relative tension, nucleus average diameter and basal contraction of pouch cells.

**SI.7 Computational model parameters.** Parameters used in model simulations in wild type and perturbed cases are listed in the SI Tables 3-6.

**SI Table 3**. Energy functions parameters used for dome shape profile formation in wild type simulation as shown in Fig. 5A-5A'. These parameters are used in the right-hand sides of the Equations 1-3.

| Energy | Definition | Interaction Type | Values | Reference |
|---|---|---|---|---|
| $E_v$ | Volume exclusion | Morse | $U_v = -V_v = 7.04\, nN.\mu m$ | Ref. [10] |
| | | | $\zeta_v = \text{v} = 0.375\ \mu m$ | |
| | | | $L_{vMax} = 0.78125\ \mu m$ | |
| $E_{nuc}$ | Size of nuclear | Morse | $U_{nuc} = 12.8\, nN.\mu m$ | This work |
| | | | $\zeta_{nuc} = 0.224\ \mu m$ | |
| | | | $V_{nuc} = 9.6\, nN.\mu m$ | |
| | | | $\gamma_{nuc} = 3.36\ \mu m$ | |
| | | | $L_{nucMax} = 3.5\ \mu m$ | |
| $E_{adhL}$ | E-cadherin | Spring | $k_{adhL} = 200\, nN/\mu m$ | Ref. [10] |
| $E_{adhB}$ | Integrin | Spring | $k_{adhB} = 40\, nN/\mu m$ | Ref. [10,13] |
| $E_{adhA}$ | Adhesion between columnar and squamous cells | Spring | $k_{adhA} = 20\, nN/\mu m$ | This work |
| | | | $L0_{adhA} = L0_{adhB} = L0_{adhL} = 0.0625\ \mu m$ | |
| $E_{cont}$ | Basal actomyosin contractility | Spring | $k_{cont} = 9\, nN/\mu m$ | Result of model calibration |
| | | | $L0_{cont} = 0.03125\ \mu m$ | |
| $E_{memb}$ | Membrane stiffness | Spring | $k_{membA} = 900\, nN/\mu m$ | Ref. [10] |
| | | | $k_{membL} = 600\, nN/\mu m$ | |
| | | | $k_{membB} = 300\, nN/\mu m$ | |
| $E_{ECM}$ | ECM stiffness | Spring | $k_{ECM} = 4500\, nN/\mu m$ | Ref. [14] |
| | | | $L0_{ECM-pouch} = -0.06\ \mu m$ | |
| | | | $L0_{ECM-BC} = -0.06\ \mu m$ | |
| | | | $L0_{ECM-perip} = 0.06\ \mu m$ | |
| $E_{vol}$ | Conserving volume of cytoplasm | Lagrange multiplier | $k_{vol} = 30\, nN/\mu m^2$ | Ref. [8] |
| | | | $\Omega_{0pouch} = \Omega_{0perip} = 65\ \mu m^2$ | |
| | | | $\Omega_{0Bc} = 20\ \mu m^2$ | |

**SI Table 4**. Parameters in the wild type simulations. Other parameters are the same as in the SI Table 3.

| Parameter | Definition | Before dome shape profile formation | After dome shape profile formation |
|---|---|---|---|
| $C_{ECM}$ | Damping coefficient (resistance to move) | $36\ nN/\mu ms$ | $36 \times 10^3\ nN/\mu ms$ |
| $E_{cont}$ | Basal actomyosin contractility | $k_{cont} = 9\ nN/\mu m$ | $k_{cont} = 0\ nN/\mu m$ |

**SI Table 5**. Parameters of energy functions representing effects of the actomyosin inhibition by using Latrunculin corresponding to Fig. 7B'. Other parameters are the same as in the last column of SI Table 4 and in SI Table 3.

| Energy | Definition | Interaction Type | Values |
|---|---|---|---|
| $E_{adhA}$ | Adhesion between columnar & squamous cells | Spring | $k_{adhA} = 0$ |

**SI Table 6**. Parameters of energy functions represent effects of ECM removal. The parameters correspond Fig. 8B'-D'. Other parameters are the same as in the last column of SI Table 4 and SI Table 3.

| Energy | Definition | Interaction Type | Values |
|---|---|---|---|
| $E_{adhB}$ | Integrin | Spring | $k_{adhB} = 0$ |
| $E_{ECM}$ | ECM stiffness | Spring | $k_{ecm} = 0$ |

**SI Table 7**. Parameters of energy functions represents effects of ECM removal and actomyosin inhibition through ROCK inhibition. The parameters correspond Figure 9B'-D'. Other parameters are the same as in the last column of SI Table 4 and SI Table 3.

| Energy | Definition | Interaction Type | Values |
|---|---|---|---|
| $E_{adhA}$ | Adhesion between columnar and squamous cells | Spring | $k_{adhA} = 0$ |
| $E_{adhB}$ | Integrin | Spring | $k_{adhB} = 0$ |
| $E_{ECM}$ | ECM stiffness | Spring | $k_{ecm} = 0$ |
